# Predicting cell-type-specific exon inclusion in the human brain reveals more complex splicing mechanisms in neurons than glia

**DOI:** 10.1101/2024.03.18.585465

**Authors:** Lieke Michielsen, Justine Hsu, Anoushka Joglekar, Natan Belchikov, Marcel J.T. Reinders, Hagen Tilgner, Ahmed Mahfouz

## Abstract

Alternative splicing contributes to molecular diversity across brain cell types. RNA-binding proteins (RBPs) regulate splicing, but the genome-wide mechanisms remain poorly understood. Here, we used RBP binding sites and/or the genomic sequence to predict exon inclusion in neurons and glia as measured by long-read single-cell data in human hippocampus and frontal cortex. We found that alternative splicing is harder to predict in neurons compared to glia in both brain regions. Comparing neurons and glia, the position of RBP binding sites in alternatively spliced exons in neurons differ more from non-variable exons indicating distinct splicing mechanisms. Model interpretation pinpointed RBPs, including QKI, potentially regulating alternative splicing between neurons and glia. Finally, using our models, we accurately predict and prioritize the effect of splicing QTLs. Taken together, our models provide new insights into the mechanisms regulating cell-type-specific alternative splicing and can accurately predict the effect of genetic variants on splicing.

## Introduction

During RNA splicing, introns are removed from the precursor mRNA. Different combinations of exons result in different mRNA isoforms, which may differ in function^1–3^. This mechanism, called alternative splicing, causes most of the complexity of human tissues and cell types; approximately 95% of all human genes are believed to be spliced in multiple ways^4,5^. Across different tissues, the brain has the highest levels of exon skipping and one of the most distinctive patterns of alternative splicing^6^.

Alternative splicing (AS) is partly regulated by RNA-binding proteins (RBPs)^7,8^, which can activate or inhibit spliceosome assembly or splice site recognition. RBFOX proteins, for instance, instruct neuronal differentiation by regulating splicing of *NIN* which in turn affects the localization of the corresponding Ninein protein^9,10^. Additionally, splicing regulation often relies on the combinatorial binding of multiple RBPs. For example, the inclusion of exon 9 of *Gabrg2* is dependent on the binding of RBFOX and NOVA^11^. Splicing simulators have taken into account splicing enhancers and silencers^12^ and a splicing code for tissue-dependent splicing has been elaborated^13–15^. However, the genome-wide mechanisms regulating splicing across different cell types remain largely unknown.

Long-read sequencing is an emerging technology that has made important contributions to RNA biology since its inception^16–20^. Long-read single-cell and single-nuclei sequencing in fresh^21,22^ and frozen^23^ tissue allows the study of alternative splicing at the cell-type level in the brain and other complex tissues. Such analyses revealed that most mouse genes show differential isoform expression across at least one pair of cell types, regions, and/or developmental time points in the brain^24,25^. In accordance with prior studies^26–28^, single-nuclei isoform RNA sequencing (SnISOr-Seq) of the human adult frontal cortex revealed that exons associated with autism spectrum disorder (ASD) are variably included across cell types^23^.

To understand (alternative) splicing mechanisms and the influence of RBPs, several computational methods have been developed. AVISPA, for instance, predicts alternative splicing in four tissues by extracting regulatory features, such as the length of the exon or the presence of RBP binding sites, from the mRNA sequence^14^. Other methods, including SpliceAI, DNABERT, Pangolin, and MTSplice, directly use the pre-mRNA sequence as input to their models^29–32^. However, none of the current methods predict cell-type-specific alternative splicing in a genome-wide manner, which is crucial for understanding splicing in heterogeneous tissues such as the brain.

Here, we present two methods to predict cell-type-specific exon inclusion using the pre-mRNA sequence and/or the presence of RBP binding sites in the hippocampus and frontal cortex. After training our machine learning models, we used model interpretation to study the mechanisms governing cell-type-specific exon inclusion. We focused on variable exons which we defined as exons for which the inclusion rates differ in neurons and glia. We found that the presence of RBP binding sites in variable exons compared to non-variable exons differs more in neurons than in glia. This indicates that the alternative splicing mechanism in neurons deviates more from the non-variable mechanism. Furthermore, we show that some RBPs, including QKI, have a big effect on exon inclusion in glia, that the regions close to the splice sites are most important for predicting exon inclusion, and that we can correctly predict and prioritize the effect of splicing QTLs and prioritize their effects.

## Results

### Predicting exon inclusion is more difficult in neurons than in glia

To define the rules governing exon inclusion in distinct cell types, we trained different models to predict cell-type-specific percent spliced-in (*Ψ*) values in the brain (Figure 1A). We focused on neurons and glia in two human brain regions, hippocampus (HPC) and frontal cortex (FC), and calculated *Ψ* values per exon by aggregating single-nuclei isoform RNA sequencing (SnISOr-Seq) reads from multiple individuals (Table 1, Methods)^23,25^. Most exons are either almost always included (*Ψ* ≈ 1) or excluded (*Ψ* ≈ 0) in an mRNA molecule (Figure 1B, S1A-C). Furthermore, most exons have similar values in neurons and glia (Figure 1C, S1D). We define exons with different inclusion rates in neurons and glia (|*ΔΨ _glia_*_−*neur*_| > 0.25) as variable exons. In HPC and FC, 2,244 and 943 exons are variable respectively (Table 1). In contrast to non-variable exons, these values show a uniform distribution of *Ψ* (Figure 1B). Even though we used a minimum of 10 reads per exon to calculate a *Ψ* value (Methods), we believe these values are reliable. When comparing the *Ψ* values of the variable exons per individual in neurons and glia, there is a clear separation between neurons and glia (Figure S2). Since most exons are almost always included, we downsampled these exons when training the models (Methods).

**Figure 1.**
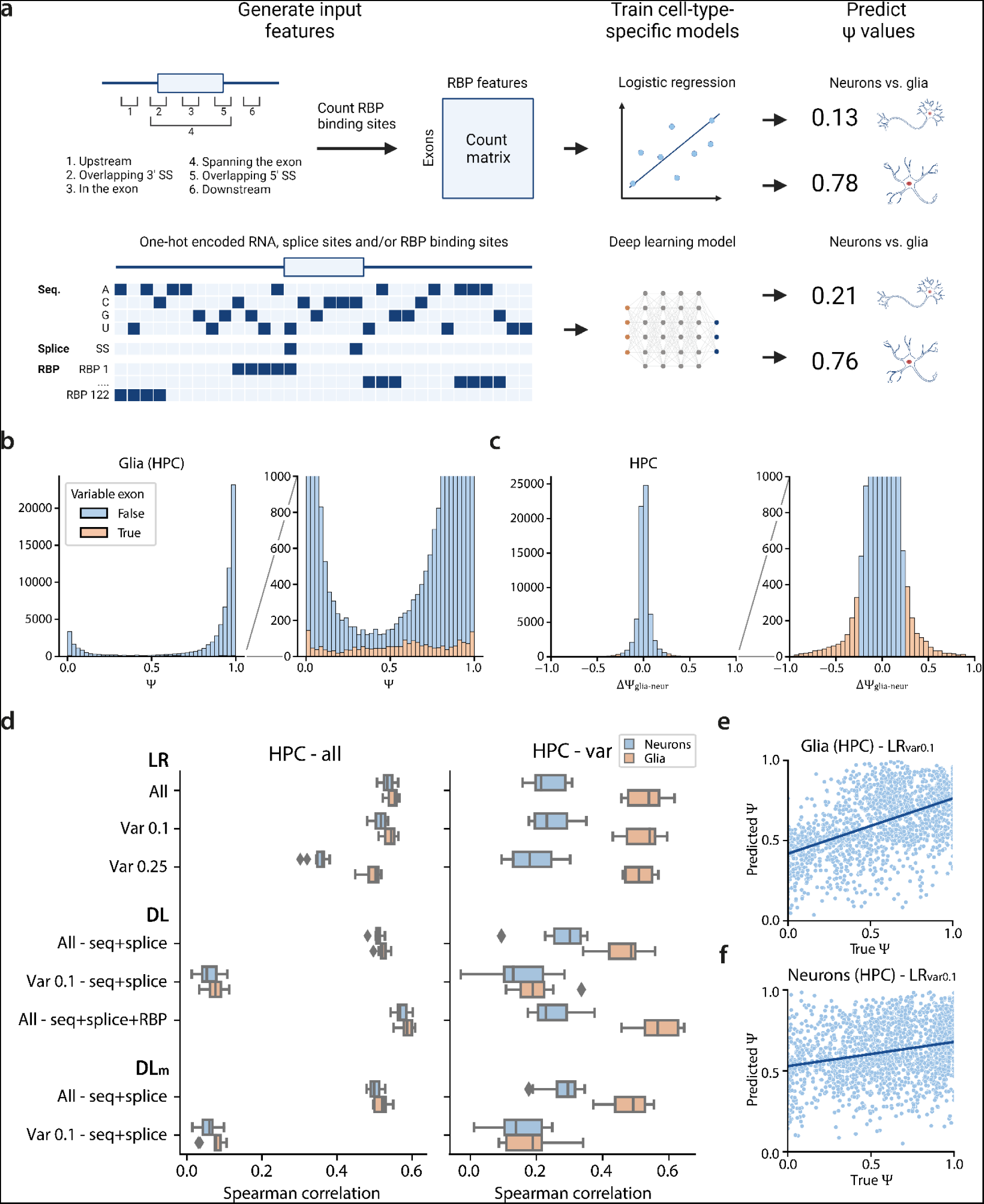
Overview and performance of the *Ψ* prediction models. **a)** Schematic overview of the models used to predict cell-type-specific *Ψ* values. **b)** Distribution of *Ψ* values of glia in the hippocampus. **c)** Distribution of *ΔΨ_glia_*_−*neur*_ for the hippocampus. **d)** Performance of the different models during 10-fold cross-validation on all exons and the variable exons in glia and neurons in the hippocampus. **e-f)** Scatterplot showing the predictions of LR_var0.1_ for variable exons in glia and neurons.

**Table 1.** The number of measured exons (exons for which at least 10 reads were sequenced in both the neurons and glia) and variable exons (|*ΔΨ _glia_*_−*neur*_| > 0.25) in the hippocampus (HPC) and frontal cortex (FC).

|  | Individuals | Measured exons | Variable exons |
| --- | --- | --- | --- |
| HPC<sup>25</sup> | 6 | 68,215 | 2,244 |
| FC<sup>23</sup> | 2 | 56,427 | 943 |

First, we used a logistic regression (LR) model to predict *Ψ* values from RBP binding sites of 122 RBPs from the ENCODE project^8^. These RBPs were measured in two cell lines (K562, HepG2), implying that this data is not brain cell-type-specific. We generated a count matrix, indicating the number of binding sites per exon for each RBP. Since the position of an RBP can influence its function^33,34^, we split these binding sites based on six possible binding locations: 1) upstream of the exon (up to 400bp), 2) overlapping the 3’ splice site, 3) in the exon, 4) spanning the exon, 5) overlapping the 5’ splice site, and 6) downstream of the exon (up to 400bp) (Figure 1A).

Any model is strongly influenced by its training data. A model trained on all exons might be dominated by the rules governing non-variable exons, while cell-type-specific inclusion effects might be overlooked. Therefore, we trained three different models using 10-fold cross-validation and either: A) all exons (LR_all_), B) exons with |*ΔΨ _glia_*_−*neur*_| > 0.1 (LR_var0.1_), or C) exons with |*ΔΨ _glia_*_−*neur*_| > 0.25 (LR_var0.25_) as training data (Table S1). When evaluating the models on all exons, LR_all_ showed the highest median Spearman correlation between true and predicted *Ψ* values on all four datasets followed by LR_var0.1_ and LR_var0.25_ (Figure 1D, S3). On hippocampal variable exons, however, LR_var0.1_ outperformed the other models (Figure 1D). The performance increase when training on variable exons indicates that the splicing mechanism in these variable exons is somewhat different from the mechanism in non-variable exons. In the frontal cortex, the performance on neurons increased when the training data became more specific, while the performance on glia decreased (Figure S3). Surprisingly, we predicted *Ψ* values more accurately in glia than neurons in both brain regions (median Spearman correlation of 0.54 vs. 0.23 in HPC, and 0.57 vs. 0.10 in FC) (Figure 1D-F, S3-4). Furthermore, using LR_var0.25_ to predict *Ψ* values of all exons resulted in lower performance for neurons compared to glia in both HPC and FC (Figure 1D, S3). Indicating that the learned splicing patterns for variable exons in neurons do not generalize to non-variable exons - likely because the underlying molecular grammar is different in the two exon sets.

### Primary sequence is more informative for neurons

The RBP binding sites used to train the logistic regression models were measured in immune and liver cancer cell lines and are thus not cell-type specific - and may reflect glial more than neuronal splicing as shown above. Furthermore, some RBPs known to be important for splicing in the brain, such as NOVA1 and NOVA2, are not included in the ENCODE data^35,36^. To test whether this caused the low performance of the models on neurons, we trained sequence-based models - which are independent of any RBP data and comparable across different cell types. We adapted the Saluki model, a hybrid convolutional and recurrent neural network that uses mRNA sequences to predict mRNA degradation rates^37^, to predict *Ψ* values (Methods) (Figure 1A, S5). The input sequence is 6,144 bp with the exon of interest centered in the middle. Since deep learning models need large training datasets, we trained a model using all exons (DL_all-seq_) and a model using exons with |*ΔΨ* _(*glia*−*neur*)_| > 0.1 (DL_var0.1-seq_).

In HPC, the LR_all_ model outperformed the DL models when evaluating performance on all exons, but on variable exons, DL_all-seq_ outperformed LR_var0.1_ for neurons (Figure 1D). For the variable exons in neurons, primary sequence is more informative than the measured ENCODE-derived RBP-binding-site data. Even though the performance increases for neurons, the performance gap between neurons and glia remains. Thus, neuronal splicing patterns probably have more complex regulation mechanisms that we do not capture with the current models. In FC, the performance of the DL models on all exons and variable exons was considerably lower compared to HPC (Figure 1D, S6). This is likely related to the size of the training data which is significantly smaller for FC than HPC (Table S1).

Next, we combined sequence and RBP binding sites by adding a channel for every RBP which indicates the presence of a binding site (DL_all-seq-RBP_) (Figure 1A, S5). This outperformed the LR models and resulted in the best-performing model for glia (median Spearman correlation of 0.54 vs. 0.57 in HPC, and 0.57 vs. 0.65 in FC) (Figure 1D, S3, S6). This improvement indicates that we can capture regulatory information from sequence beyond those present in RBP data alone. For neurons, however, DL_all-seq-RBP_ had lower performance than DL_all-seq_, again confirming that the ENCODE RBP data is more informative for glia than neurons.

We also trained DL models that do not use splice sites or only use RBPs as input for the neurons and glia in HPC to understand how the input channels affect performance (Figure S7). Omitting splice sites only slightly decreased the performance, which indicates that the model can recognize the splice sites quite easily from the sequence itself. For glia, using the RBPs as the only input feature results in a comparable performance to the LR_all_ model (median Spearman correlation of 0.55 vs. 0.54) and an even better performance than sequence and splice sites only (median Spearman correlation of 0.49). However, for neurons, we observe the opposite; using RBP binding sites reduces performance compared to the DL_all-seq_ model (median Spearman correlation of 0.23 vs. 0.30).

### Exon inclusion mechanisms are conserved between human and mouse

As cell-type-specific alternative splicing is partially conserved between humans and mice^25^, we hypothesized that adding mouse data to our model would increase performance. We combined human HPC data with mouse HPC^25^. Since mouse FC data is not available, we combined human FC with data from the mouse visual cortex (VIS). While these two cortical regions are not identical, they do share many common characteristics. Especially in mouse HPC, few exons are variable (528) compared to VIS (1,404) (Table S2, Figure S8). Although DL_all-seq-RBP_ performed best in glia, we only trained models with sequence and splice sites as input channels (DL_all-seq-m_, DL_var01-seq-m_) since RBP binding sites were not measured in mouse cell lines. In HPC, the performance on variable exons of both cell types slightly increased by adding the mouse data (Figure 1D). On FC, the performance on all exons increased as well (Figure S6), supporting our hypothesis that not enough training data was available to train these models on human exons alone. Similar to the human data, glial *Ψ* values were easier to predict than neuronal ones in mice (Figure S9).

### The splicing mechanisms in neurons diverged more than in glia

Our above results show that neuronal *Ψ* values are harder to predict than glial regardless of model or input data. Hence, splicing mechanisms in neurons might be different than in glia and more complex. However, *Ψ* values could be biased, making it easier to predict in glia. To exclude the latter, we used the hippocampus data to assess whether glia and neurons are similar in terms of 1) *Ψ*-value distributions, 2) heterogeneity within each cell type, and 3) variation across individuals.

First, comparing *Ψ* distributions, more values are close to 0 or 1 in glia than neurons (Figure S10AB), which is most apparent for the non-variable exons (two-sided Kolmogorov-Smirnov test, p-value < 2.2e-16). For variable exons (Figure S10B), however, both distributions are not different (two-sided Kolmogorov-Smirnov test, p-value = 0.44). Thus, data distribution differences cannot explain all observed differences between neurons and glia.

Second, to quantify the heterogeneity within a cell type, we measured the difference in *Ψ* values between finer cell-type classifications. For neurons, we compared the inhibitory and excitatory neurons, and for glia, we compared oligodendrocytes and astrocytes. Within glia, we have more variable exons (|*ΔΨ*| > 0.25) compared to neurons (831 vs. 745). In neurons, more exons have an extreme difference (|*ΔΨ*| > 0.5) (92 vs. 70) (Figure S10CD). Compared to the total exon number defined for both cell types in neurons and glia (28,296 and 27,047 respectively), both numbers are small. Thus, this cannot explain the difference in performance between neurons and glia.

Third, to compare the variance across individuals for glia and, separately, for neurons, we calculated *Ψ* values per individual instead of using the aggregated counts. We calculated the variance for an exon only if ≥3 individuals have ≥10 reads for that exon in both neurons and glia. For both non-variable and variable exons, the variance is higher in glia (two-sided paired Wilcoxon signed-rank test, p-value = 1.3e-28 and 8.9e-5 respectively) (Figure S10E). Thus, the data do not explain observed differences in performance between neurons and glia.

We then hypothesized that splicing mechanisms regulating variable exons in neurons might differ from the non-variable exons. To test this hypothesis, we compared the RBP binding profiles between variable and non-variable exons in neurons and glia (Figure 2A). We performed these comparisons for exons with a high (≥ 0.5) and a low *Ψ* value (< 0.5) separately. The binding profiles between variable and non-variable exons differ significantly more in neurons compared to glia in HPC (Figure 2B) and FC (Figure 2C). Non-variable exons with high *Ψ* values more often have a binding site at the 3’ splice site for splicing factors such as U2AF1, U2AF2, and SF3B4 compared to non-variable exons with low *Ψ* values (Figure 2D, S11AB). In glia, variable exons show a similar pattern (Figure 2E, S11AB). However, binding sites for these splicing factors cannot differentiate between exons with high and low *Ψ* values in neurons (Figure 2F, S11AB), indicating that these RBP binding sites are likely not used in neurons. In the hippocampus, PTBP1 differs the most between neurons and glia (Figure S11C). PTBP1 is a position-dependent RBP: binding within or upstream of an exon represses splicing while binding downstream activates splicing in HeLa cells^38^. Our RBP binding profiles contradict these known mechanisms. In HeLa cells, however, PTBP1 is highly and PTBP2 is lowly expressed, while this is vice versa in the hippocampus (Figure S12). PTBP1 RBP binding profiles obtained from non-brain cell lines are thus less likely to reflect splicing mechanisms in the hippocampus. Strikingly, the binding profile of PTBP1 in variable exons in neurons is again considerably different from the variable exons in glia and the non-variable exons. There is no position-dependent regulation and no difference between exons with a high and low *Ψ*. In the hippocampus, only one RBP, HNRNPC, showed the opposite pattern with larger differences in glia compared to neurons (Figure S11D).

**Figure 2.**
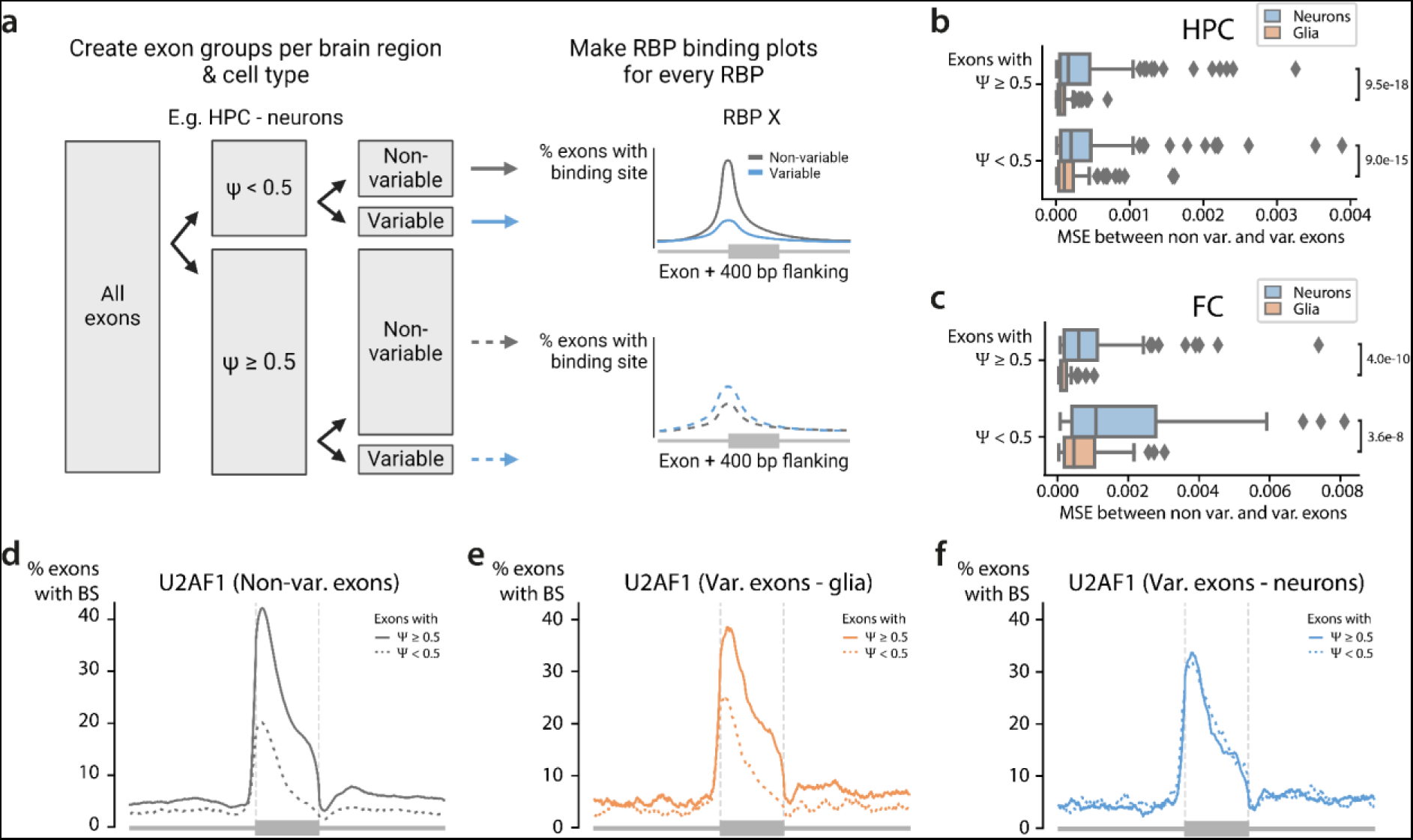
The difference in RBP binding profiles between non-variable and variable exons. **a)** Schematic overview showing how to generate the RBP binding profiles of non-variable (|*ΔΨ _glia_*_−*neur*_| ≤ 0.25) and variable (|*ΔΨ _glia_*_−*neur*_| > 0.25) exons in neurons in the hippocampus. We generated these RBP binding profiles for every RBP and split the exons based on their *Ψ* value (threshold = 0.5) and their variability We calculated the mean-squared error (MSE) between the profiles in non-variable and variable exons. We do this for the exons with a high and low *Ψ* value resulting in four comparisons per RBP. **b-c)** Boxplot showing the MSE between the RBP profiles in non-variable and variable exons in neurons (blue) and non-variable and variable exons in glia (orange) for the **b)** hippocampus and **c)** frontal cortex. Every point in the boxplot is one RBP. P-values are calculated using a two-sided paired Wilcoxon signed-rank test. **d-f)** Binding profile of U2AF1 in **d)** non-variable exons, **e)** variable exons in glia, and **f)** variable exons in neurons.

### Interpretation of LR models reveals cell-type-specific splicing mechanisms

To further pinpoint the factors underlying differences in splicing between glia and neurons, we analyzed the coefficients of the logistic regression models. These coefficients reflect the importance of each RBP binding position in regulating cell-type specific splicing. We compared the coefficients of four models for the hippocampus (two cell types, and two training sets) and focused on features present in at least 50 exons and with a coefficient > 0.05 in at least one model (191 out of 732 features). The model coefficients first cluster based on which exons are used during training (all vs. variable) (Figure 3A). This clustering indicates that the mechanisms for non-variable and variable exons, represented by the LR_all_ and LR_var0.1_, differ more than the cell-type-specific mechanisms. The RBPs cluster into two groups: features with positive and features with negative coefficients (Figure 3A). As expected, splicing repressors, which are part of the heterogeneous nuclear ribonucleoproteins (hnRNP) family^39^, have a largely negative weight in all models (Figure 3B). PTBP1, for which we saw a difference between the non-variable and variable exons in the hippocampus, is a member of the hnRNP family and has a potential position-dependent effect in glia based on the RBP binding profiles (Figure S11C). The LR_var0.1-glia-HPC_ model correctly learned this effect: PTBP1 binding at the 3’ splice site and within the exon have coefficients of −0.05 and 0.01 respectively. The model coefficient for PTBP1 binding at the 3’ splice site is among the ten features that differ the most between glia and neurons (Figure 3C, LR_var0.1-glia-HPC_ vs LR_var0.1-neur-HPC_) which indicates a potential cell-type-specific effect corresponding to the established switch between PTBP1 and PTBP2^40–42^.

**Figure 3.**
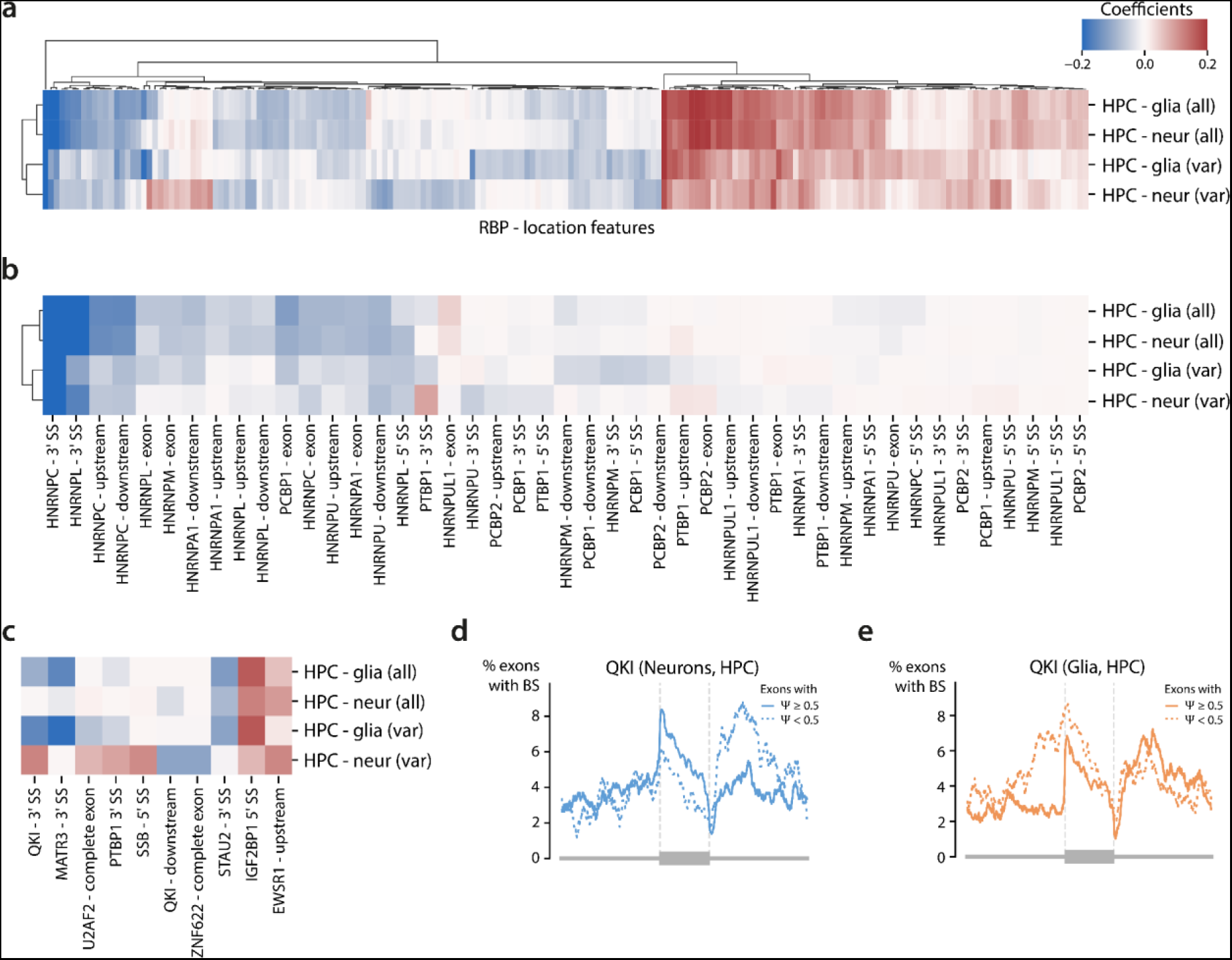
Interpretation of the logistic regression models. **a)** Heatmap showing the coefficients for the RBP-location features in the different logistic regression models. The input features are filtered using a minimum of 50 RBP sites and a value of at least 0.05 in one of the models. The values are clipped between −0.2 and 0.2. **b)** Heatmap showing coefficients of hnRNPs in the different models. **c)** Heatmap showing the top 10 cell-type-specific input features with the biggest difference between HPC - glia (var) and HPC - neur (var). **d-e)** Binding profiles of QKI in variable exons in neurons and glia.

QKI binding at the 3’ splice site has the strongest cell-type-specific effect in the hippocampus (model coefficient = −0.15 vs. 0.12 for glia and neurons respectively), which reflects differences in the RBP binding profiles (Figure 3D-E). In glia, a binding site that overlaps the 3’ splice site leads to lower inclusion rates, while the opposite happens in neurons. In the scRNA-seq data, QKI has higher expression in glia compared to neurons in the hippocampus (Wilcoxon rank sum test, adj. p-value < 2.2e-16) (Figure S13). Both observations correspond to the known mechanism of QKI, which inhibits splicing by competing with the core splicing machinery^10,43^. In mice, QKI is important during myelination and oligodendrocyte differentiation^44,45^. Its role in the human brain is less studied, but a role in oligodendrocyte formation and Schizophrenia has been suggested^46,47^. Interestingly, variable exons are enriched for QKI binding sites compared to non-variable exons (Fisher’s exact test, adj. p-value = 1.6e-13). Besides the 3’ splice site, QKI binding downstream of an exon is also in the top 10 cell-type-specific features. The effect of QKI downstream of an exon is the opposite compared to QKI binding at the 3’ splice site, which indicates a potential position-dependent effect of QKI. Such position-dependent regulation of QKI has been shown in lung cancer^48^ but, to our knowledge, not in the brain.

In contrast to QKI, most of the cell-type-specific RBPs identified using our LR models are neither differentially expressed nor differentially spliced. Exceptions are STAU2, which is upregulated in neurons (Wilcoxon rank sum test, adj. p-value < 3.39e-16), and EWSR1, which is differentially spliced (Table S3). The latter could indicate that distinct isoforms of EWSR1 influence RNA splicing in different ways.

### The sequence close to the splice sites is most important for predicting exon inclusion

Given that the RBP-binding-site data is not brain-specific and that it lacked measurements from some key RBPs, we set out to identify sequence features that influence *Ψ* predictions in the deep learning models. We used *in-silico* saturation mutagenesis (ISM, Methods) to systematically predict how nucleotide substitutions in the input sequence affect the predicted *Ψ* value^49–52^. Since DL_var0.1_ performed considerably worse than DL_all_ (Figure 1D), we focused on interpreting DL_all_ for glia in the hippocampus, which had higher prediction accuracy than neurons, instead of looking for cell-type-specific effects.

Since ISM is computationally expensive, we mutated the input sequence of the 9,929 exons with |*ΔΨ* _(*glia*−*neur*)_| > 0.1 instead of all exons. The ISM score indicates how much a mutation increases or decreases the predicted *Ψ* value compared to the average prediction at that position for that sequence (Methods). As expected, mutations around the splice sites and within the exon strongly affect the predicted *Ψ* value (Figure 4A). These results reflect the known importance of the splice site’s consensus sequence to be recognized by the splicing machinery. The two nucleotides before and after the exon - the AG acceptor and GU donor dinucleotides - have the strongest predicted effects. Looking at the maximum absolute ISM score, only mutations within a range of 50bp upstream of the 3’ splice site and 150bp downstream of the 5’ splice site have a value > 0.1 (Figure S14). This is in line with a recent computational model that predicted human splice sites using a window of 400bp on each side of the splice site and obtained an overall accuracy of 96%^53^. However, smaller values of >0.05 could be observed across almost the whole input sequence. Although distant splicing regulators have been reported^54^, potential variability in distant motifs and/or their position may prevent their detection by our model.

**Figure 4.**
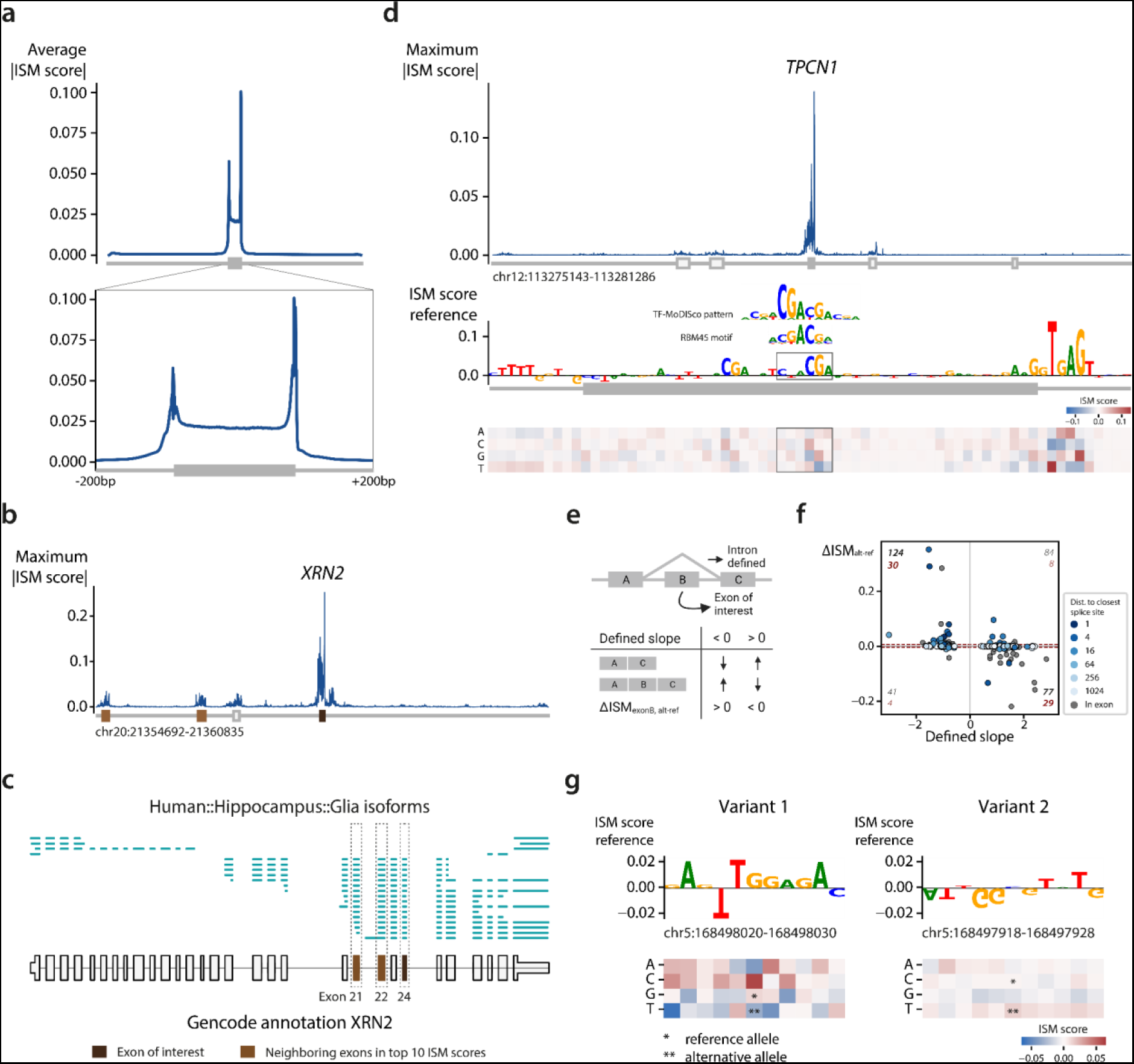
Interpretation of the deep learning model for glia in the hippocampus. **a)** Average absolute ISM score across the 9,929 exons. The mutations within the exons are binned in 300 bins. The zoomed-in plot ranges from 200bp upstream of the 3’ splice site to 200bp downstream of the 5’ splice site. **b)** Mutation profile for an exon in *XRN2.* The colors of the exons below the profile indicate the exon of interest and the neighboring exons which have an ISM score in the top 10. **c)** Single-cell long reads for *XRN2.* Each line is a single cDNA molecule. The bottom black track shows the Gencode annotation. **d)** Mutation profile for an exon in *TPCN1*. In the exon, a motif corresponding to RBM45 is found. **e)** Schematic overview of the sQTL analysis. **f)** Scatterplot showing the predicted effect for each variant. The color of the points indicates the distance to the closest splice site. A grey dot means that a variant falls within the exon of interest. The numbers in black and red indicate the number of predictions in a quadrant when no threshold and a threshold of 0.005 are used respectively. **g)** ISM scores for two variants related to the same exon of *RARS1.* A negative effect, corresponding to the positive slope, is predicted for the first variant. A smaller, but positive effect is predicted for the second variant.

Besides the region around the exon of interest, we observed higher-than-average ISM scores within nearby exons and their flanking region (Figure S15). The enrichment of RBP binding sites in these regions could explain the higher scores. Alternatively, our model potentially recognized coordinated events between exons. To test this, we selected the top 10 exons with the highest absolute ISM scores within their neighboring exons and visualized the single-cell long reads from our data that span both exons (see Methods). These reads can inform whether the two exons pair non-randomly (thus in coordination^21,23,55,56^) or randomly. Exon 24 in *XRN2* appeared twice in the top 10 list with two neighboring exons (exons 21 and 22) strongly influencing its *Ψ* value (Table S4). All three exons (21, 22, and 24) have a *Ψ* value of around 0.8 and the exons are either all included or all excluded in the single-cell long-read data, suggesting these exons are mutually associated (Figure 4BC). Mutations affecting the inclusion of one of these exons will most likely affect the other exons as well. In the top 10 scores, four other cases could pinpoint exon coordination events (Figure S16-19). In the remaining four cases, the exons pair randomly, so there is no evidence of exon-exon coordination (Figure S20-23).

To further interpret sequences with a high ISM score, we used TF-MoDISco^57^ to identify motifs in sequences with large effects on exon inclusion. Since the region around the splice site had the highest ISM scores, many of the top motifs identified by TF-MoDISco correspond to the consensus splice sites and associated motifs, including the well-known AG acceptor dinucleotide, the poly-pyrimidine tract (PPT) upstream of the exon, and the extended splice donor motif with the GU dinucleotide (Supplementary File 1, Figure S24). We also found motifs that match known RBP binding motifs, which were not in our input data for the LR model, and hence could not be tested for cell-type-specific effects. For example, we found a motif corresponding to RBM45 in exon 12 of *TPCN1* (Figure 4D, Table S4), which seems to promote exon inclusion. RBM45 regulates constitutive splicing and can probably activate or repress the inclusion of an exon, but the exact mechanisms are currently unknown^58^. Taken together, characterizing important sequence features from DL models can identify splicing regulators beyond those we can identify based on available RBP measurements.

### Prioritizing the effect of splice QTLs using the DL models

So far, we showed how LR and DL model interpretations can be used to reveal the regulatory mechanisms of RBP governing cell-type-specific exon inclusion. Besides this fundamental knowledge, we can use our DL models to predict the effects of genetic variants on splicing. Accurately predicting these effects can help prioritize variants of interest. To test the relevance of our model predictions for genetic variants, we used splicing quantitative trait loci (sQTLs) from the hippocampus data from GTEx v8^59^. Variants in this dataset are linked to intron excision ratios instead of exon inclusion. We extracted introns and their corresponding variant(s) that span an exon in our data and predicted the effect of the variant(s) on that exon (Figure 4E). In total, 326 variants are within the input range of our model. These variants correspond to 122 introns and 158 exons. Some introns thus span multiple exons and most introns have multiple variants linked. For every variant, a slope indicates whether the corresponding intron is excised more or less compared to the reference allele. We expect negative slopes to correspond to an increased *Ψ* value of the exon of interest which would result in *ΔISM* _(*alt*−*ref*)_ > 0. Conversely a positive slope would result in *ΔISM* _(*alt*−*ref*)_ < 0 (Figure 4E). However, more complex scenarios, such as a variant affecting adjacent exons, may arise as well.

Using our model, we predicted an effect (|*ΔISM* _(*alt*−*ref*)_| > 0.005) for 71 out of 326 variants which corresponds to 61 of the 122 introns. For 83% (59 out of 71) of these variants, our model predicts the expected effect correctly (Figure 4F, S25). Most of the variants with an effect are very close to the splice sites: 74.6% are within the exon or a distance of 15bp to either the 3’ or 5’ splice site. These cases thus affect most likely exonic splicing enhancers or the binding of U1 and U2 snRNA. For 14 of 61 introns where our model did not predict an effect, all corresponding variants are outside of the intron itself. Here, the splicing of adjacent exons is most likely altered instead of our exon of interest. For 2 of these 14 exons, all variants are even outside of the gene itself.

Three exons have multiple corresponding variants with a predicted effect. For exon 15 in *ZNF880* (Table S4), three variants have a predicted expected effect. The other two exons, however, have two variants with a contradicting predicted effect. In both cases, the variant with the biggest predicted effect is in line with the slope of the sQTL of the intron. For exon 25 in *RARS1* (Table S4), for instance, variant one is located in the exon (168,498,025; G → T) and variant two is located before the exon (168,497,923; C → T). For variant one, our model predicted the expected effect, while our model predicted the opposite for variant two (Figure 4G). Variant one, the variant with the biggest and correctly predicted effect, is located in a binding site for SRSF1 according to eCLIP data^8^. RNA recognition motif 2 (RRM2) of SRSF1 interacts with the GGA motif. A G → T mutation in the first nucleotide will thus hinder the binding of SRSF1^60^. Variant two is located in a stretch of G’s. At this location, there’s a binding site for ELAVL1, a protein regulating mRNA stability, and hnRNP family member HNRNPK, which tends to repress splicing^8^. Using the DL models, we can thus correctly predict the effect for most sQTLs and prioritize their effects.

## Discussion

We trained logistic regression and deep learning models to predict cell-type-specific exon inclusion in human brain samples. Since this is the first attempt to leverage long-read single-cell sequencing data for this task, we can use our models to decipher the grammar underlying cell-type specificity of splicing. Using model interpretation, we pinpointed interesting RBPs, such as QKI, that could drive differential splicing between neurons and glia. Furthermore, we show that the location of RBP binding sites differs more between variable and non-variable exons in neurons compared to glia. This indicates that the splicing mechanisms controlling exon inclusion in neurons are more different compared to the general mechanism.

For most RBPs, RBP binding profiles of non-variable exons with high and low *Ψ* values showed distinct patterns. Considering U2AF1 for example, exons with a high *Ψ* value are more likely to have a binding site close to the 3’ splice site compared to exons with a low *Ψ* value. These RBPs behave differently in variable exons in neurons, and for most RBPs the difference between exons with a low and high *Ψ* value is missing. These features are thus not informative for neurons, which explains the low performance of the logistic regression models on neurons. The U2AF heterodimer, composed of U2AF1 and U2AF2, is believed to bind every polypyrimidine tract and AG dinucleotide in 3’ splice site regions^61–63^. Binding may not happen on specific sites repressed by other factors. The potential binding sites are still there, but they might be used by a competing RBP in neurons. Interestingly, most RBPs are not differentially expressed or differentially spliced between neurons and glia. For these RBPs, post-translational modifications, such as phosphorylation, might differ between neurons and glia and could change their function^64,65^. Furthermore, RBP binding sites measured in non-brain cell lines might not always be representative of splicing in the hippocampus and frontal cortex. The expression of RBPs can differ dramatically between the non-brain and brain tissues as was seen for PTBP1.

The deep learning models, however, also perform poorly on the variable exons in neurons. The model trained on all exons focuses only on learning the general splicing mechanisms, and the model trained on the variable exons might not have enough training data. In glia, the model trained on all exons performs well on the variable exons. Again indicating that the variable exons in glia follow the rules of the general splicing mechanisms more. The worse performance of the DL_all-seq_ models on neurons, in combination with the distinct RBP binding profiles, supports our conclusion that the splicing mechanisms in variable exons in neurons diverged from the mechanisms in non-variable exons.

A potential explanation, in line with the diverged RBP binding sites, is that splicing in neurons is less sequence-dependent. Other factors, such as chromatin features and polymerase speed^66–79^, RNA methylation^80–82^ as well as other modifications, and transcription factor binding sites^83^, influence splicing as well. These features might explain the difference between neurons and glia. Altered chromatin accessibility or RNA methylation, could, for instance, explain why certain RBP binding sites are not used in neurons anymore. Furthermore, neuronal genes - by definition more expressed in neurons - are more susceptible to missplicing^84^. While we did not focus on missplicing, this indicates that splicing mechanisms might be different in neurons. Also, the gene expression of human neurons diverged faster from other primates compared to glia^85^. A similar divergence could have occurred with the splicing mechanisms.

For the deep learning model, we tested the effect of different lengths for the input sequence. Even though all lengths showed a very similar performance, we used a relatively long input sequence (6,144 bp) which had the advantage that we could predict the effect of more mutations. When predicting the effect of sQTLs, however, we predict a strong effect mainly for variants close to the exon of interest. The region close to the splice sites, however, still contributes the most to the predictions. This is in contrast to splice site predictions from SpliceAI, for which an input sequence of 10kb significantly outperforms 400 bp^29^. SPLAM, however, outperforms SpliceAI while only using 400 bp^53^. Of note, this does not preclude the mechanistic influence on splicing decisions by motifs further upstream. Rather, these data suggest that such distant RNA binding sites are highly variable regarding their position to the exon. This variability in position could prevent the model from detecting such motifs. Similar observations have been made for models that predict gene expression. Even though the best-performing model uses a long input sequence (196kb), only one-third of the receptive field is used during predictions and distal enhancers are not captured by the model^51,86^.

Another possible advantage of a longer input sequence is that it would be possible to look at coordinated events. Exons in the human brain are often mutually associated or mutually exclusive^23,55,87–89^. Such events can even be cell-type-specific. For instance, two neighboring exons in *WDR49* are perfectly coordinated in astrocytes only^23^. Using our model, the ISM scores within neighboring exons are higher than the ISM scores of the rest of the sequence. For some exons, these higher scores indeed indicate that there is exon-exon coordination. Since exon-exon coordination is so common, predicting such events might be more beneficial than focusing on individual exons.

Furthermore, the longer input sequence enables predicting the effect of more splicing QTLs. However, most variants the model predicted an effect for are near the splice sites. For these variants, the model obtained a high accuracy (83%) and could be used to prioritize the effect of splicing QTLs as well. Nonetheless, a limitation of the current DL models is that they lack cell-type specificity. The DL models need substantial training data, so training on all exons yielded the highest performance. As a consequence, these models focused on the general splicing mechanisms and yielded better performance on variable exons in glia than neurons.

In conclusion, to increase our understanding of (alternative) splicing in the brain, we trained two types of models to predict exon inclusion in neurons and glia of the hippocampus and frontal cortex. Ideally, these models make perfect predictions such that they can be used in the clinic for predicting the effects of variants or for personal splicing predictions. The performance of our models, however, is not optimal yet. Nevertheless, we show how model interpretation yields important biological discoveries including the different mechanisms in neurons and glia. This demonstrates the potential of using long-read single-cell data for this task.

## Methods

### Calculating cell-type-specific Ψ values

For the human data, we combined SnISOr-Seq data from 6 individuals for the hippocampus and 2 individuals for the frontal cortex (Table 1). For the mouse data, we combined ScISOr-Seq2 data from two mice for the hippocampus and two mice for the visual cortex (Table S2). Scisorseqr was used to map and align reads to GRCh38 for human and mm10 for mouse to identify splice sites for each dataset separately^24^. We used IsoQuant to correct the splice sites^90^. Using all exons appearing as an internal exon in a read, we calculated:

- The number of long-read molecules containing this exon (both splice sites included): *X_in_*
- The number of long-read molecules assigned to the same gene as the exon, which skipped the exon but includes ≥50 bases on both sides: *X_out_*
- The number of long-read molecules supporting the acceptor of the exon and ending on the exon: *X_acc In_*
- The number of long-read molecules supporting the donor of the exon and ending on the exon: *X_don In_*
- The number of long-read molecules overlapping the exon: *X_tot_*

Non-annotated exons with one or two annotated splice sites, ≥70 bases of non-exonic (in the annotation) bases, were excluded as intron-retention events or alternative acceptors/donors. We then calculated

- 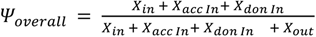
- 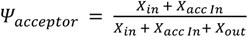
- 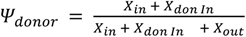

If

- 0.02 ≤ *Ψ_i_* ≤ 0.98 where *i* ∈ {*overall,acceptor,donor*} in the pseudo-bulk data, the exon was kept.

Next, we filtered exons based on the number of reads. We only calculate *Ψ_overall_* for a cell type in a certain brain region if at least 10 long-read UMIs are sequenced across the different individuals (*X_tot_* ≥ 10). Since individuals of different datasets were sequenced using a different read depth, we normalized the read counts by dividing it by the total number of reads for an individual before calculating *Ψ_overall_*. We then calculated *Ψ_overall_* for each cell type (*Ψ_neur_* and *Ψ_glia_*) for the hippocampus and frontal cortex. If there were not enough reads, *Ψ_overall_* for that exon and cell type was set to “NA”. We used the cell-type labels defined in the original datasets. For neurons, we grouped the inhibitory and excitatory neurons. For glia, we grouped the oligodendrocytes, astrocytes, and oligodendrocyte precursor cells.

### Downsampling cell-type-specific Ψ values

In the human data, many exons (30,273 out of 68,215 for the hippocampus and 45,680 out of 56,427 for the frontal cortex) have *Ψ_neur_* > 0.9, *Ψ_glia_* > 0.9, and *ΔΨ* _(*glia*−*neur*)_ < 0.03. We downsampled these to 5,000 to make the distribution less skewed towards one.

In the mouse hippocampus data, 18,351 out of 23,857 exons have *Ψ* = 1 in neurons and glia, so we downsampled these to 5,000 as well. For the visual cortex, 27,073 out of 48,515 exons have *Ψ_neur_* > 0.9, *Ψ_glia_* > 0.9 and *ΔΨ* _(*glia*−*neur*)_ < 0.03. We downsampled these to 5,000.

### RBP-binding-site data

We downloaded the eCLIP data for 122 RBPs from the ENCODE portal (https://www.encodeproject.org/metadata/?status=released&internal_tags=ENCORE&assay_title=eCLIP&biosample_ontology.term_name=K562&target.investigated_as=RNA+binding+ protein&biosample_ontology.term_name=HepG2&assembly=GRCh38&type=Experiment&fil es.processed=true). From this file list, we used the BED files that store the peaks per replicate. We merged peaks from different replicates or cell lines to ensure one BED file per RBP.

### Logistic regression models

The logistic regression model is implemented as one fully connected layer between the input features (the RBP binding sites) and the output (the *Ψ* value) with a sigmoid activation function to scale the output between 0 and 1. The models are single-task models which means that a separate model was trained for each cell type.

When training the model, we use a binary cross entropy loss with L1 and L2 regularization (alpha = 0.001, and L1-ratio = 0.7), a learning rate of 0.005, and a batch size of 256.

As input for the logistic regression models, we counted the number of peaks in the BED files for every RBP and exon at six locations: 1) upstream of the exon (maximum 400 bp away from the splice site), 2) overlapping the 3’ splice site, 3) within the exon, 4) spanning the exon, 5) overlapping the 5 splice site, and 6) downstream of the exon (maximum 400 bp away from the splice site). Since we used the eCLIP data of 122 RBPs and there are 6 possible locations, this resulted in 732 input features for every exon (Figure 1A). If peaks of different replicates were overlapping, we counted those peaks only once.

The logistic regression model is implemented in PyTorch Lightning^91,92^.

### Deep learning models

We adapted the architecture of the Saluki model^37^ by removing one convolutional layer, shortening the maximum sequence from 12,288 to 6,144 bp, and changing the final activation function to a sigmoid activation function (Figure S5). The exon of interest was centered in the middle of the input sequence. The input channels of the model depend on the input features used (sequence, splice sites, and/or RBP binding sites). For the sequence, we one-hot encoded the sequence which results in four channels. If the splice sites were used as input, this added an extra channel that indicates the start and end of the exon of interest. When adding the RBP binding sites, we add a channel for every RBP which one-hot encodes whether there is a binding site in any of the replicates of the eCLIP data for that RBP based on the BED files.

Similar to the logistic regression models, we trained a model for every cell type separately. Even though we adapted the Saluki model, we retrained all the weights in the model. When adding the mouse data, we adapted the same approach as Saluki and made the model a multi-head model where the weights of the convolutional and recurrent neuronal network layers are shared and the weights of the fully connected layer are species-specific (Figure S5).

When training the model, we used the same hyperparameters, including the learning rate, batch size, etc., as the original Saluki model (Figure S5).

For the hippocampus, we tested how input-sequence length and the number of convolutional layers affect the performance. The benefit of a longer input sequence is that the model can learn how long-distance interactions of regulatory elements affect splicing, but these models contain more parameters and are more difficult to train. The different models performed similarly which indicates that the most important information is close to the splice sites of the exon (Figure S26). The model using 6,144 bp and five channels performed slightly better for both neurons and glia and therefore we used it during all the experiments.

### Evaluation

We evaluated the performance of the models using a 10-fold cross-validation. We ensured that the same set of exons was always in the same test fold such that we could compare the performance of the models. Exons from the same gene were always in the same test fold. When training the deep learning models on human and mouse data simultaneously, we ensured that human-mouse homologs were in the same test fold. We used biomart to obtain the human-mouse homologs.

Some exons do not have any binding sites measured for any of the RBPs (5,560 exons in the hippocampus and 3,462 in the frontal cortex). This could for instance happen if certain genes were not expressed in the cell lines when the RBP binding sites were measured. Since the logistic regression model could not predict a *Ψ* value for these exons, we filtered these from the training set used for the logistic regression model and from all test sets (to enable a fair comparison between the logistic regression and deep learning models). The deep learning models are thus trained on more exons (Table S1). In the test set, there are 1,827 and 1,072 variable exons for the hippocampus and frontal cortex respectively.

We trained all models five times for every fold and averaged the predictions across these five runs. We evaluated the performance by calculating the Spearman correlation between the true and predicted *Ψ* values.

### RBP binding profiles

We generated RBP binding profiles by calculating the fraction of exons with an RBP binding site at every location (400 bp upstream of the exon −400 bp downstream of the exon). Since exons have variable lengths, we bin the exons in 50 bins and only include exons that are at least 50 bp long in the analysis. We also filter out exons without RBP binding sites.

We calculate these profiles for four different groups of exons: 1) non-variable exons with *Ψ* ≥ 0.5, 2) non-variable exons with *Ψ* < 0.5, 3) variable exons with *Ψ* ≥ 0.5, and 4) variable exons with *Ψ* < 0.5.

To define how much the mechanisms in the variable exons diverged from non-variable exons, we calculate the mean-squared error between the RBP binding profiles of the non-variable and variable exons. We do this for the exons with a high and low *Ψ* separately.

### RBP expression data

We used the 10X scRNA-seq data from the same samples to look at the gene expression of the RBPs that were measured using the eCLIP data. We used Seurat v4 for the analysis^93^. To create the heatmap in Figure S13, we normalized the data per dataset using log normalization and a scale factor of 1e6. Next, we averaged the expression over all the cells. For the frontal cortex, we next averaged over the three datasets. We plotted the *log* (*x* + 1) values.

We used the FindConservedMarkers() function using the default parameters (including Bonferroni multiple testing correction) from Seurat to find differentially expressed RBPs between neurons and glia. This tests for differentially expressed genes per individual and merges the results.

### Interpretation of logistic regression model

For the interpretation of the logistic regression models, we looked at the coefficients of the input features. To obtain one value per input feature, we average the coefficients of the 10 folds and 5 runs per fold (so the average across 50 models in total).

We only compared the coefficients across models, if there were at least 50 exons with a binding site for that input feature.

### *In-silico* saturation mutagenesis

We used *in-silico* saturation mutagenesis (ISM) to interpret how nucleotide substitutions in the input sequence affect the predictions. We did this for 9,929 exons using the DL_all-seq-m_ model trained on glia in the hippocampus. For every exon, we used the fold for which that exon was in the test set. We averaged the predictions across the 5 runs. The ISM score is defined as follows: 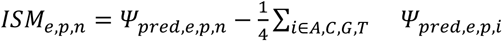 where *e* is the exon we predict the *Ψ* value for, and *p* and *n* are the position and nucleotide used at that position respectively. To visualize the ISM scores across the input sequence, we binned the upstream region, exon, and downstream region since they all had varying lengths.

### Analysis of neighboring exons

We compared the ISM scores at the exon of interest, the neighboring exons, and the remaining sequence. We extracted the locations of annotated exons from GENCODE v35^94^. The ISM scores for the exon of interest and the neighboring exons include the flanking sequence of 150 bp upstream and downstream of the exon.

Next, we selected ten exons on the positive strand with the highest absolute ISM scores in a neighboring exon. We visualized the long-reads spanning both exons using ScisorWiz^95^

### Motif discovery

We used TF-MoDISco-lite (v2.2.0)^57^ to discover motifs using the ISM scores as input. When creating the report, we compare the found motifs to the position weight matrices from oRNAment which includes motifs found using RNAcompete and RNA-bind-n-seq experiments^8,96,97^.

TF-MoDISco-lite is designed for DNA instead of RNA and tries both the forward strand and its reverse complement when finding seqlets (parts of the sequence with high ISM scores). We used the results file, to check whether the forward or reverse complement was used to generate a motif. We kept forward motifs if at least for 25 sequences the forward strand was used. We kept the reverse motif if at least for 25 sequences the reverse complement was used.

### sQTL analysis

We used the sQTLs defined for the hippocampus in GTEx v8. These variants are linked to introns instead of exons. We predicted the effect for variants that are linked to an intron that spans an exon in our dataset (Figure 4E). For most introns, there are multiple variants linked to them. We only predicted the effect for the best variants (the variants with the lowest p-value for an intron). For most introns, there were still more than two after this filter.

### Exon naming

We named exons after their position in the transcript by counting their position in the GTF file. A conversion from exon names to genomic coordinates can be found in Table S4.

## Code and data availability

The *Ψ* values, predictions, and RBP binding profiles are available on Zenodo: https://zenodo.org/doi/10.5281/zenodo.10669666.

The code to reproduce the figures, and train your logistic regression or deep learning models can be found on GitHub: https://github.com/lcmmichielsen/PSI_pred.

## Acknowledgements

Figures 1A, 2A, 4E, and S5 were created with BioRender.com.

