## Supplement for "Predicting cell-type-specific exon inclusion in the human brain reveals more complex splicing mechanisms in neurons than glia"

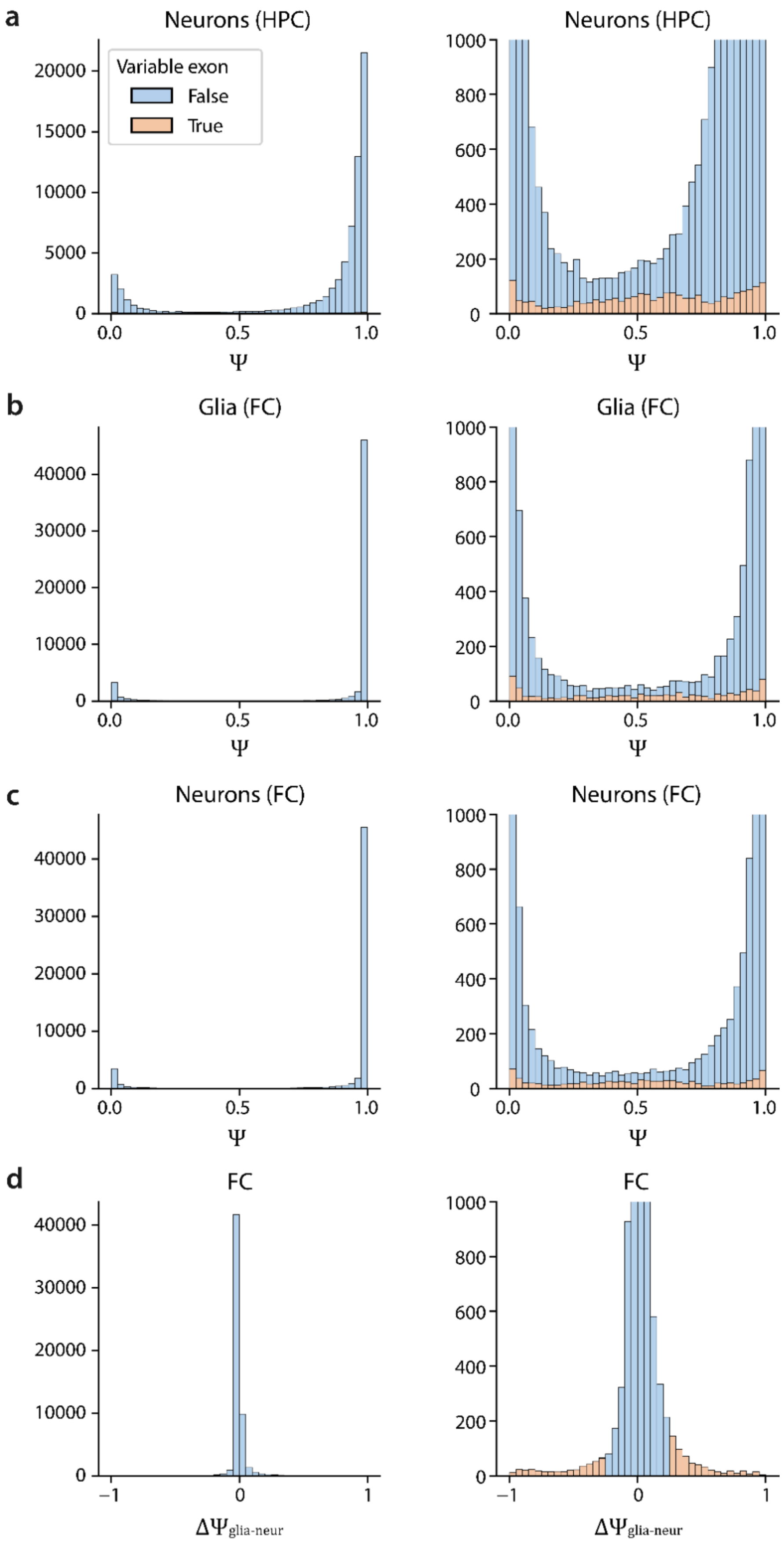


**Figure S1** Distribution of **a)** $\Psi$ of neurons in the hippocampus, **b)** $\Psi$ of glia in the frontal cortex, **c)** $\Psi$ of neurons in the frontal cortex, **d)** $\Delta\Psi_{glia-neur}$ in the frontal cortex

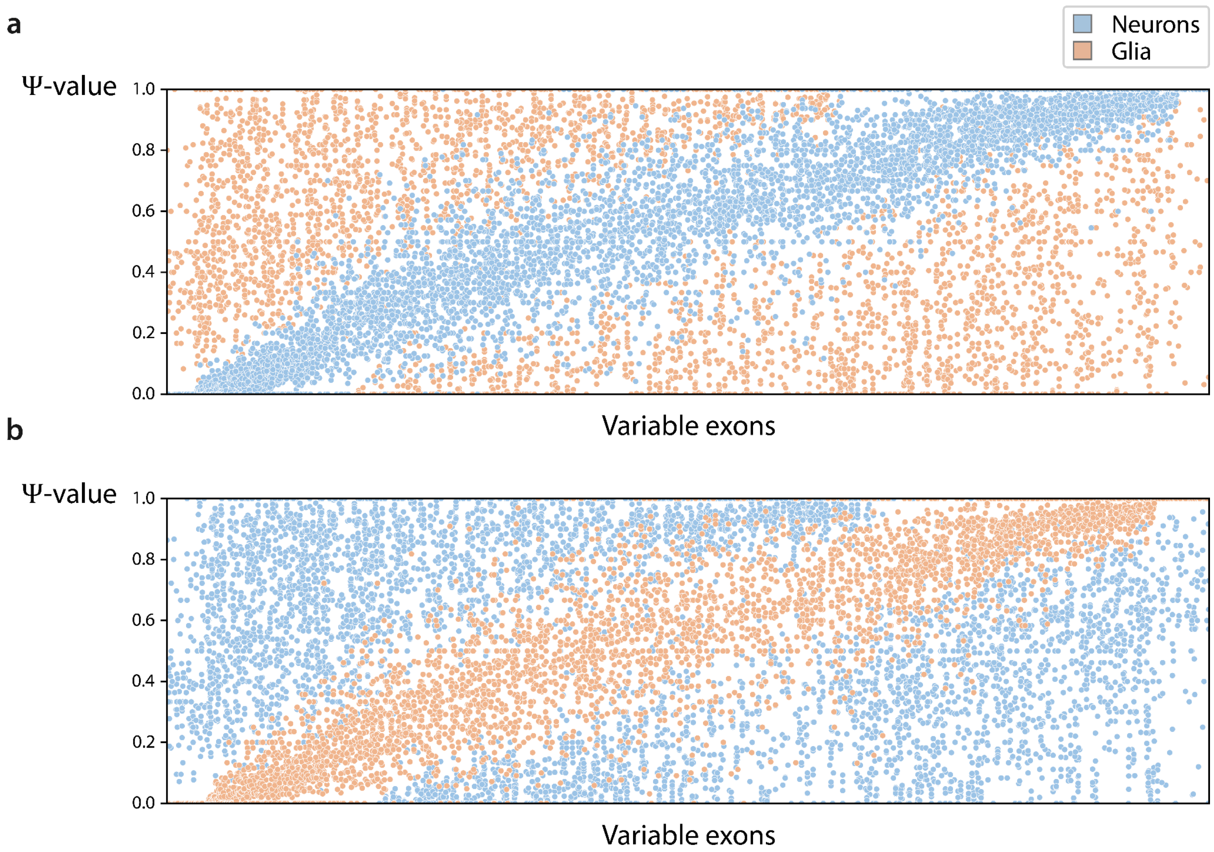


**Figure S2** Ψ values per individual for the variable exons in neurons and glia. Every column in the plot represents one variable exon. The variable exons are sorted based on the average Ψ value in **a)** neurons and **b)** glia.

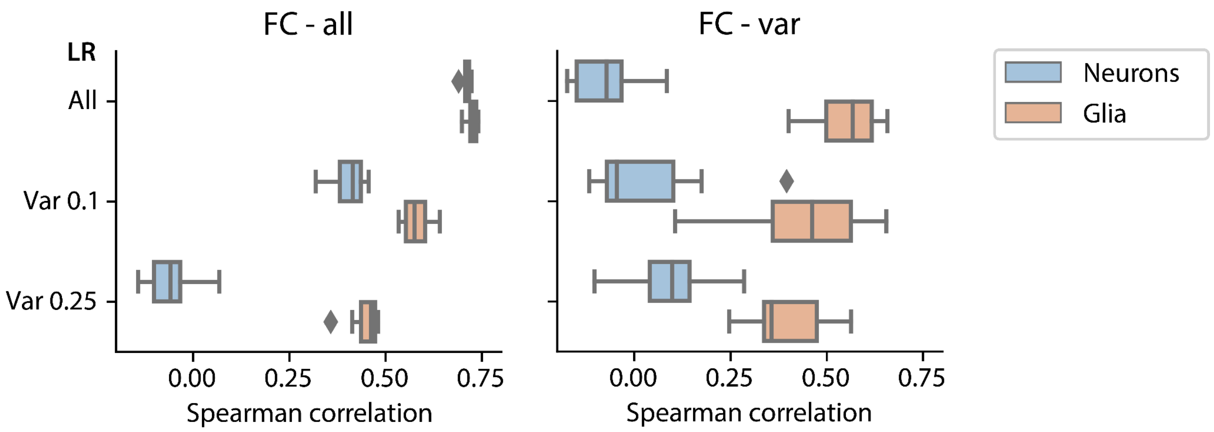


**Figure S3** Performance of the different logistic regression models during 10-fold cross-validation on all exons and the variable exons in glia and neurons in the frontal cortex.

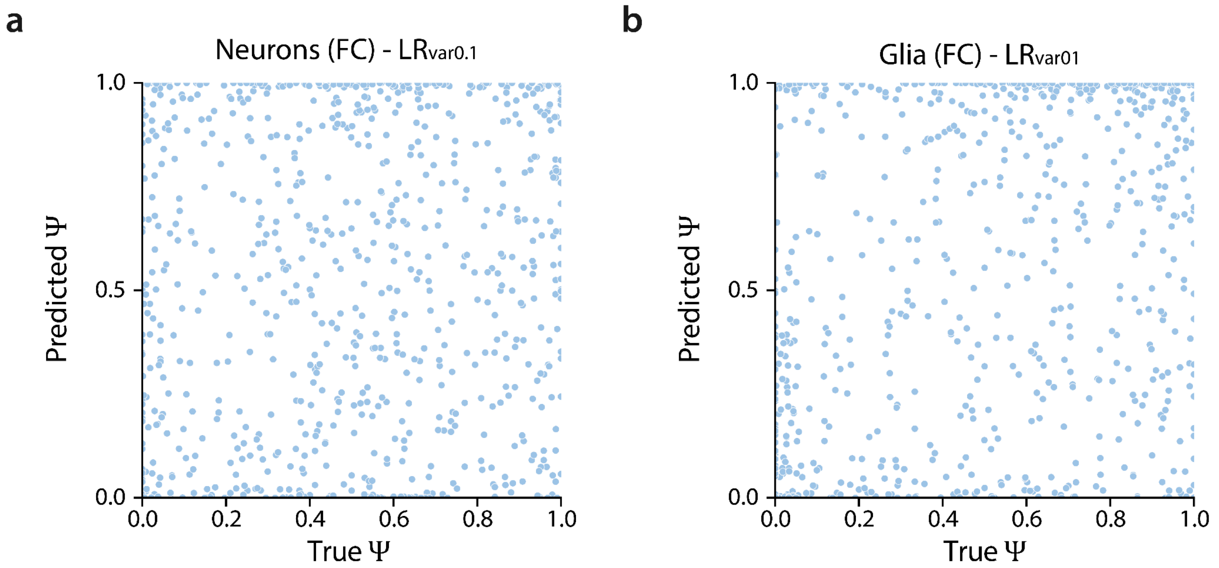


**Figure S4** Scatterplot and kernel density plot showing the predictions of the LR models on the neurons and glia in the frontal cortex on the variable exons only.

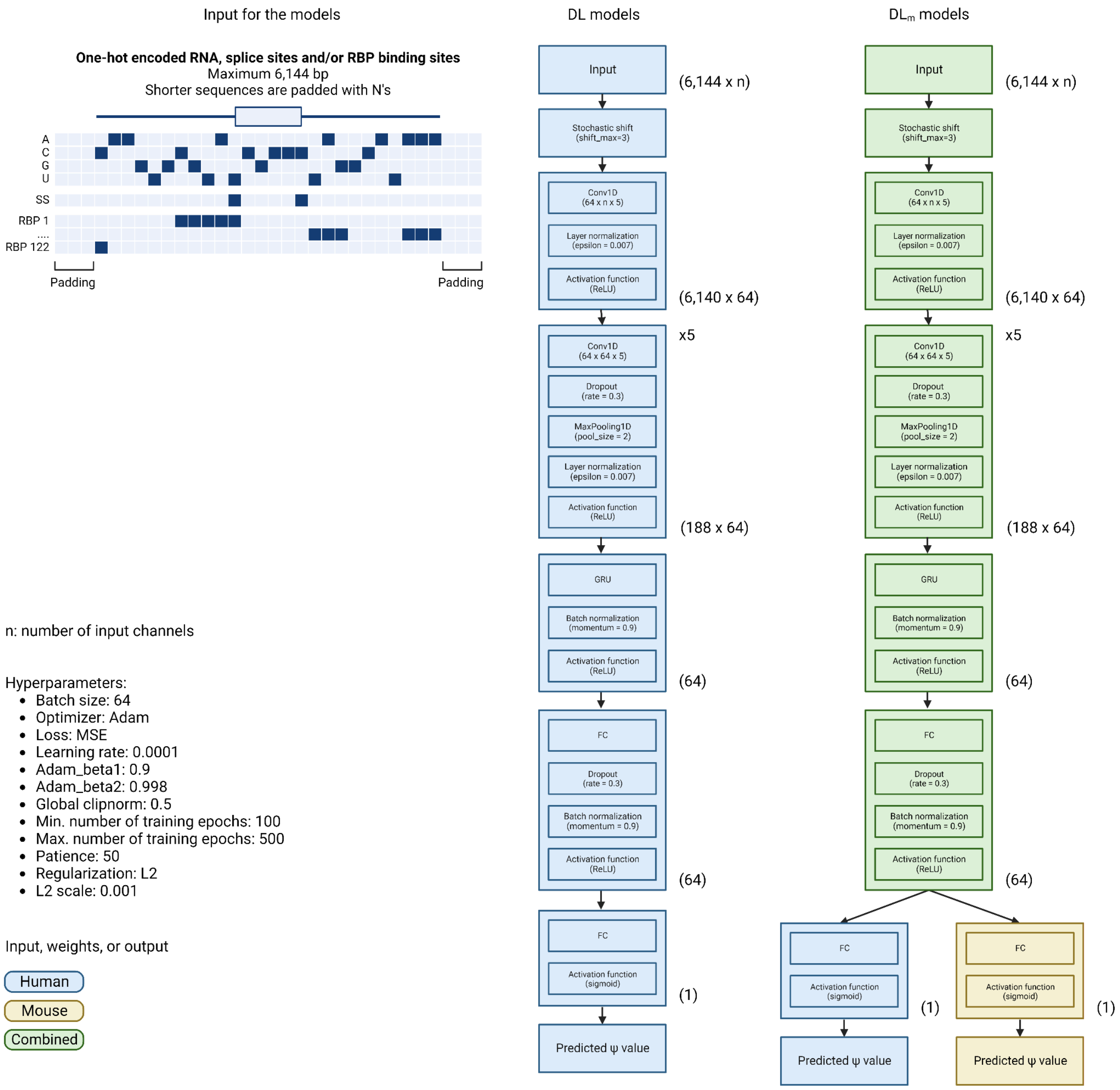


**Figure S5** Architecture of the DL and $DL_m$ models. The tuples on the right side of the blocks indicate the output shape.

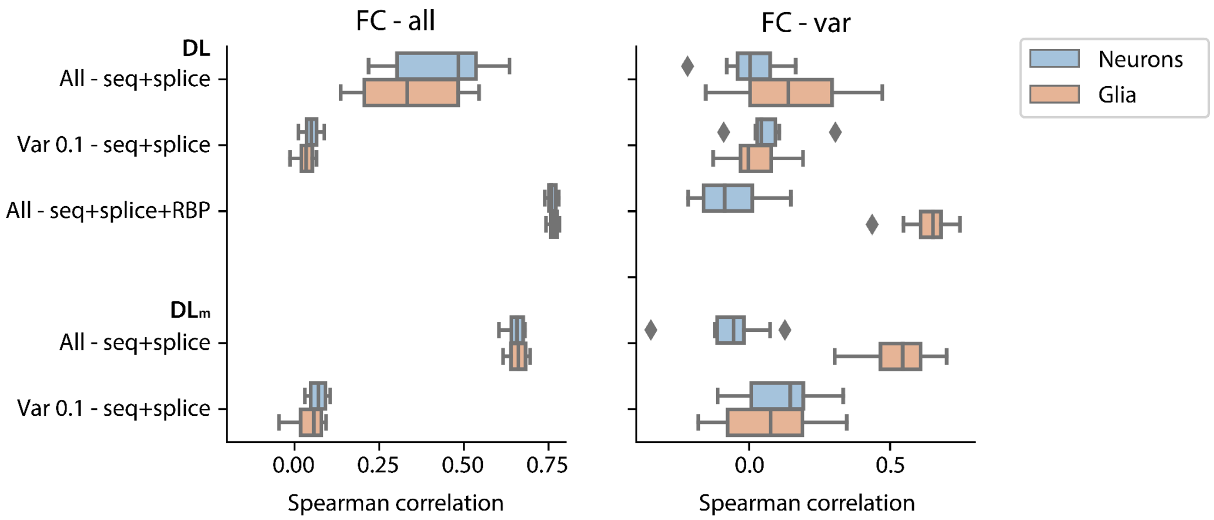


**Figure S6** Performance of the different deep learning models during 10-fold cross-validation on all exons and the variable exons in glia and neurons in the frontal cortex.

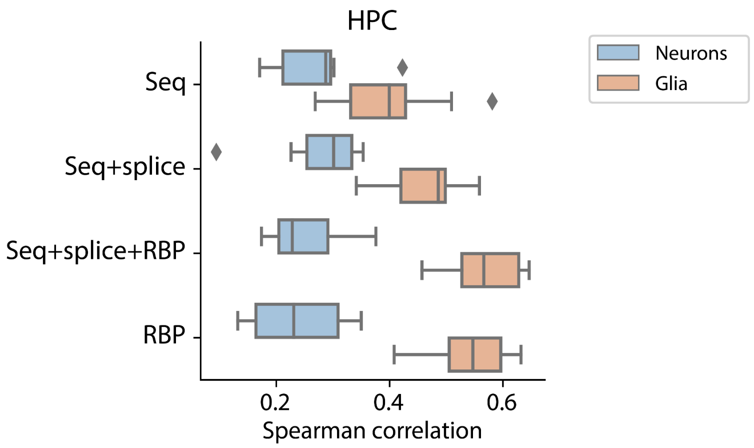


**Figure S7** Performance of the $DL_{all}$ models during the 10-fold cross-validation on neurons and glia in the HPC when trained using different input features.

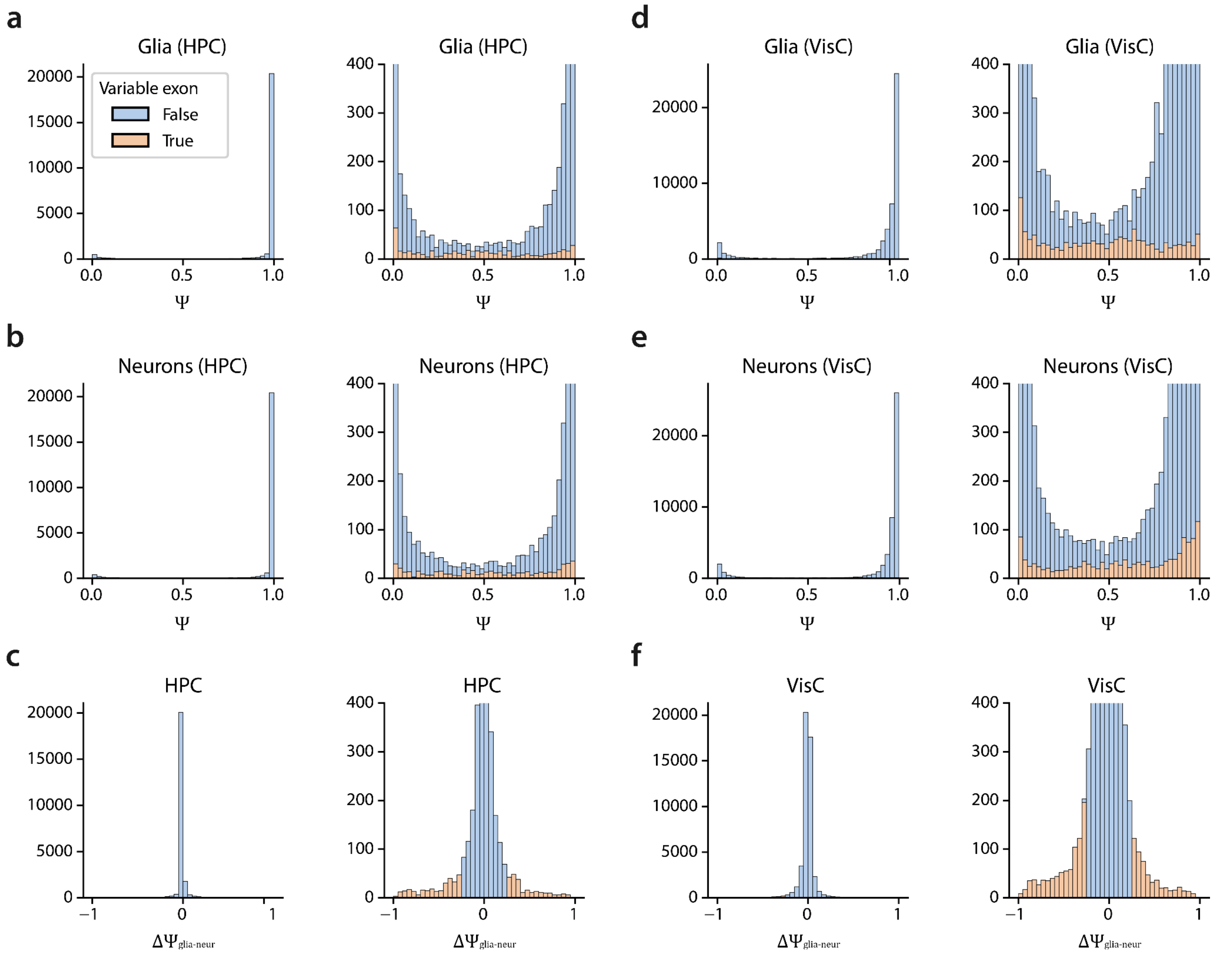


**Figure S8** Distribution of $\Psi$ of **a,d)** glia and **b,e)** neurons in the **a-b)** hippocampus and **d-e)** visual cortex in mouse, **c,f)** Distribution of $\Delta\Psi_{glia-neur}$ in the hippocampus and visual cortex in mouse.

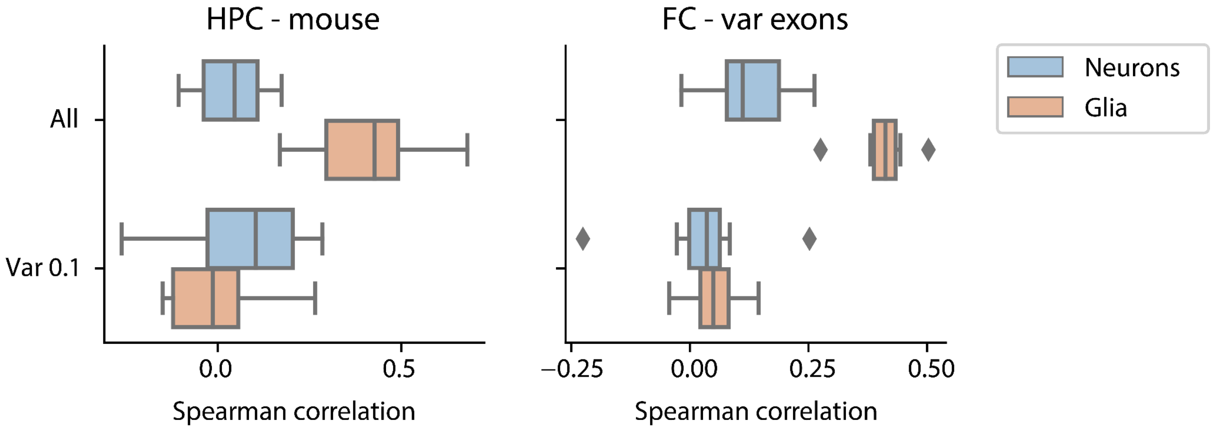


**Figure S9** Performance of $DL_{all\text{-}seq\text{-}m}$ and $DL_{var01\text{-}seq\text{-}m}$ during the 10-fold cross-validation on variable mouse exons from neurons and glia in the HPC and visual cortex.

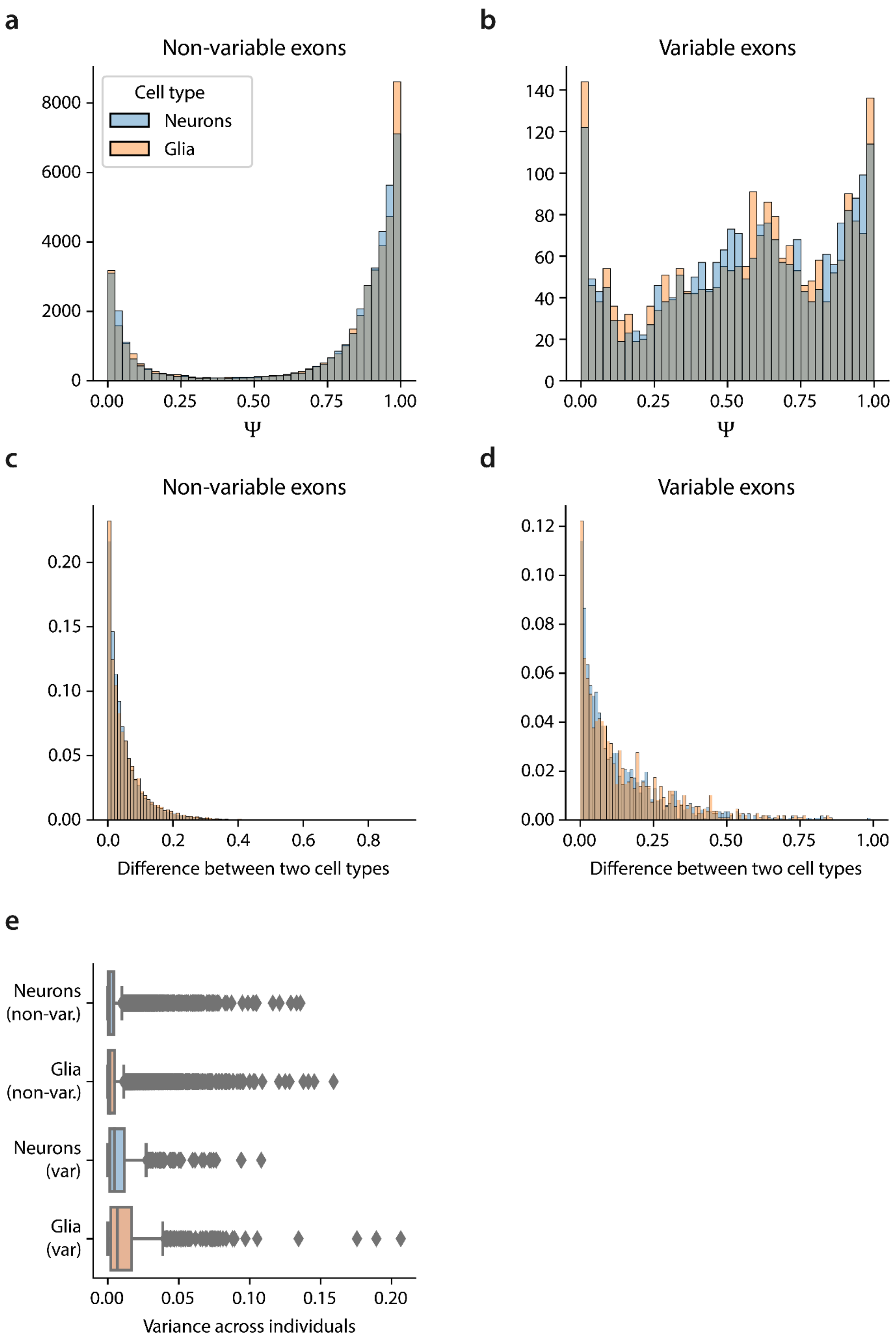


**Figure S10 a-b)** Distribution of $\Psi$ split for non-variable and variable exons, **c-d)** $|\Psi_{celltype1-celltype2}|$ for the non-variable and variable exons. For neurons, inhibitory and excitatory neurons are compared. For glia, oligodendrocytes and astrocytes are compared. **e)** Variance of $\Psi$ across the individuals.

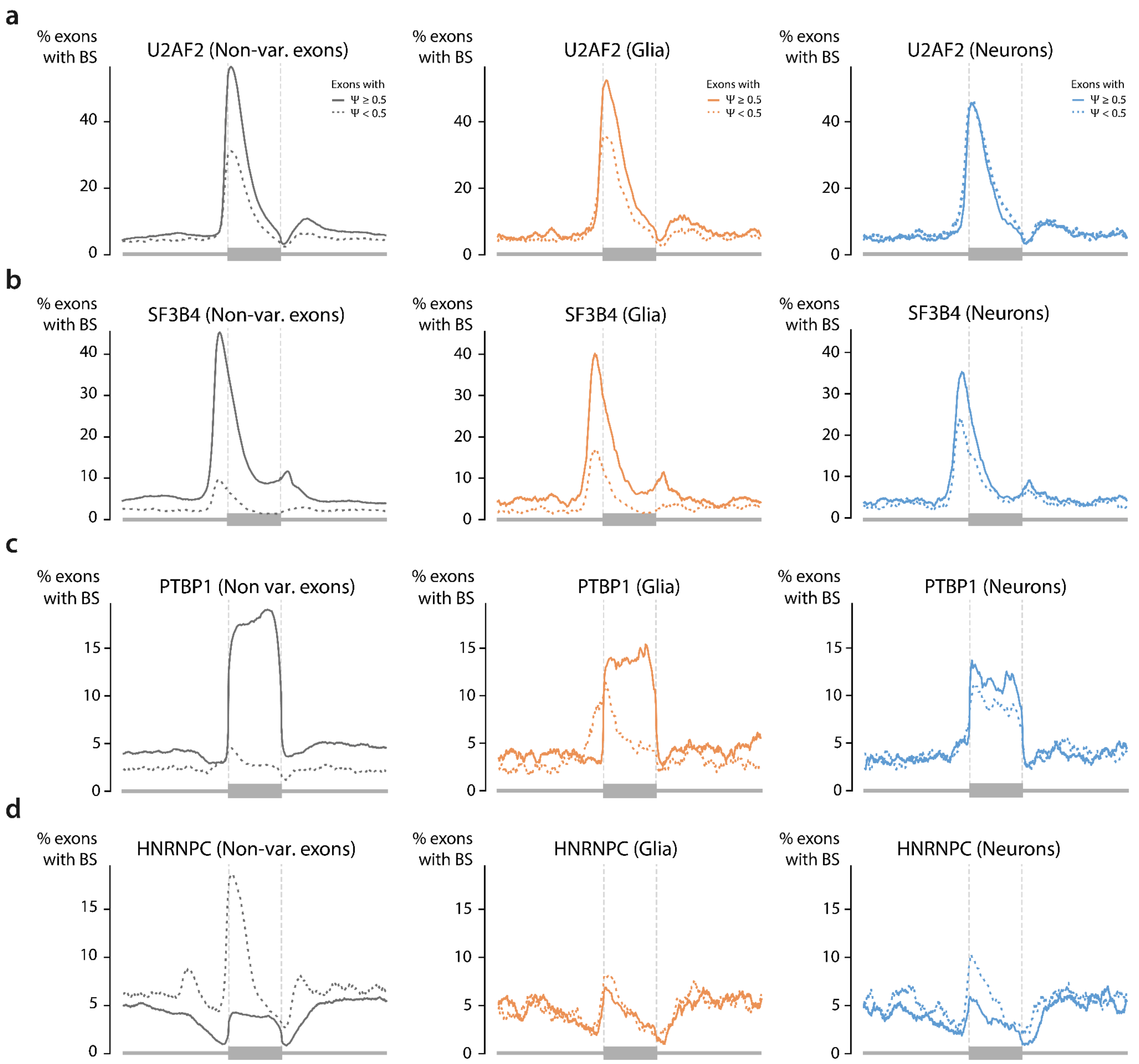


**Figure S11** RBP binding profile of **a)** U2AF2, **b)** SF3B4, **c)** PTBP1, and **d)** HNRNPC in non-variable, variable exons in glia, and variable exons in neurons in the hippocampus.

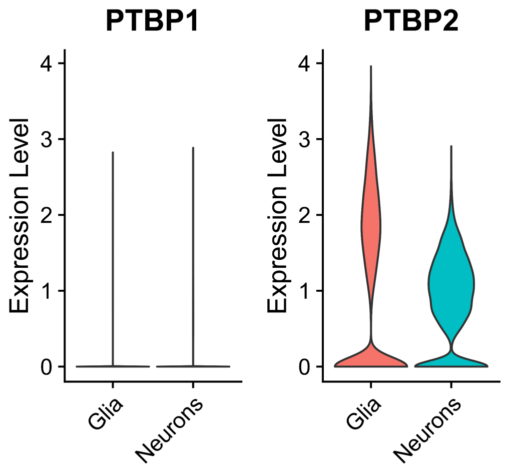


**Figure S12** Expression of PTBP1 and PTBP2 in glia and neurons in the scRNA-seq data of the hippocampus.

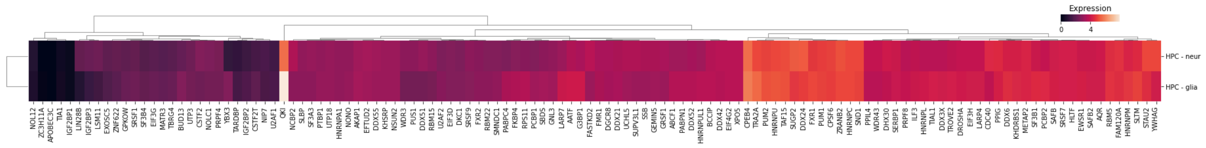


**Figure S13** Expression of RBPs in the scRNA-seq data of the hippocampus.

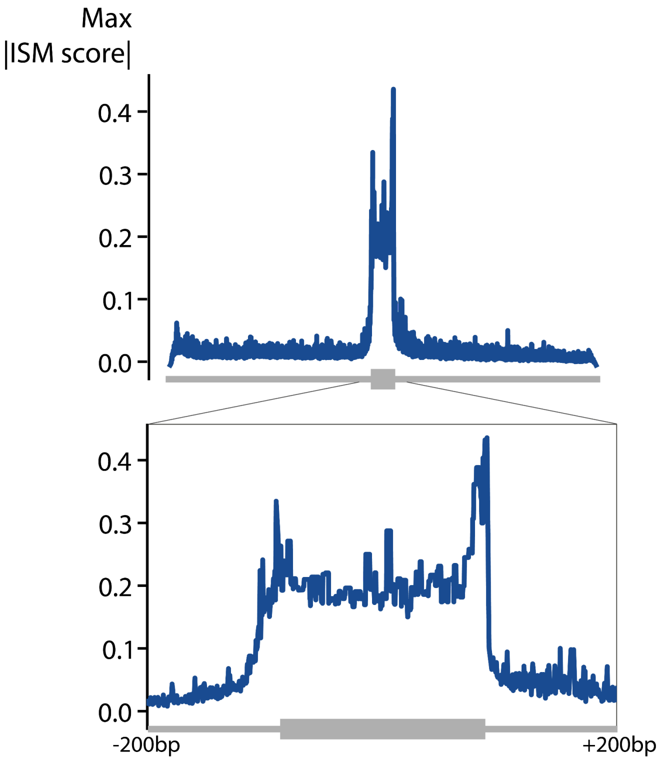


**Figure S14** Maximum absolute ISM score over all sequences. Values above 0.1 are only seen in the range of 50bp upstream of the 3’ splice site until 150 downstream of the 5’ splice site. The zoomed-in plot ranges from 200bp upstream of the 3’ splice site to 200bp downstream of the 5’ splice site.

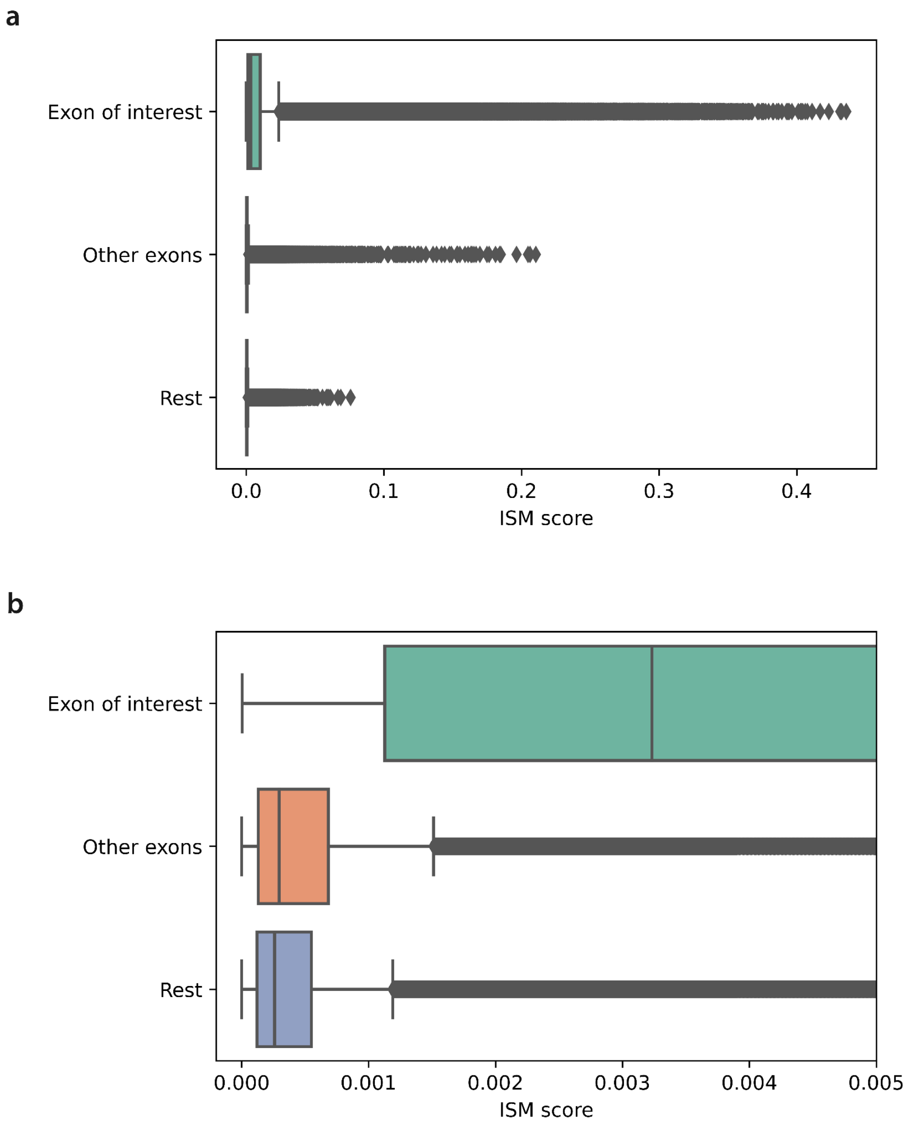


**Figure S15** Boxplot showing the maximum absolute ISM score of each position. Positions are grouped based on whether they fall in the exon of interest, another exon, or the remaining sequence. **a)** Complete boxplot, **b)** zoomed in boxplot to show the difference between the other exons and the remaining sequence.

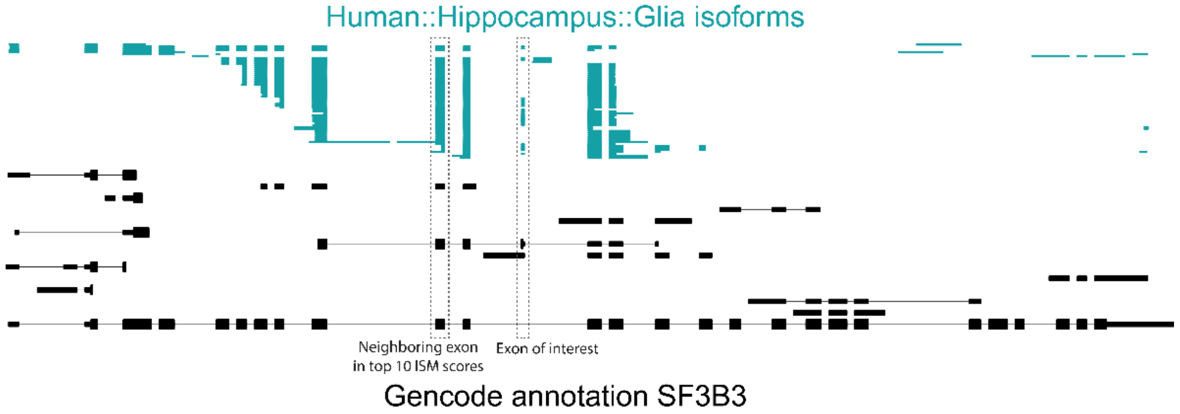


**Figure S16** Potential coordination in SF3B3. If the neighboring exon is not included, the exon of interest is also not included. A mutation in the neighboring exon that decreases its Ψ value, could thus decrease the Ψ of the exon of interest as well.

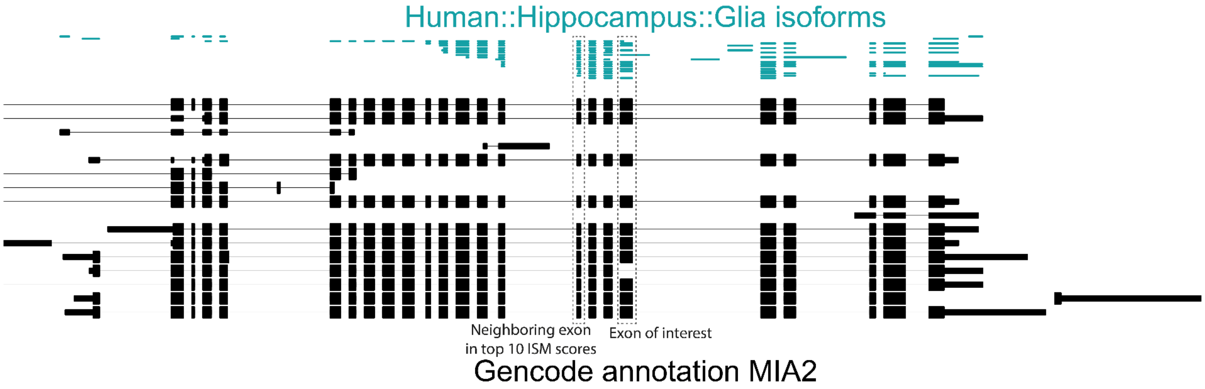


**Figure S17** Potential coordination of MIA2. If the neighboring exon is included, the exon of interest is also included. A mutation in the neighboring exon that decreases its Ψ value, could thus decrease the Ψ of the exon of interest as well.

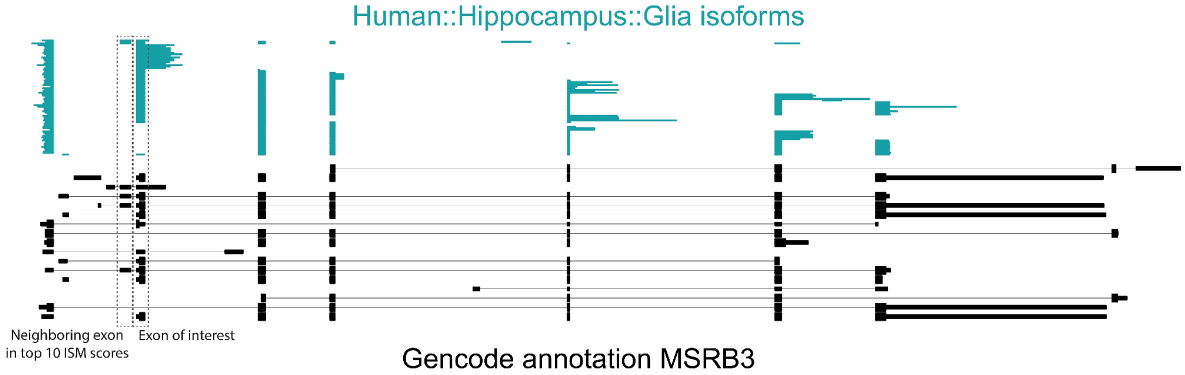


**Figure S18** Potential coordination in MSRB3. The Ψ value of the neighboring exon is low, but if this exon is included, the exon of interest is also included. Especially a mutation that increases the inclusion of the neighboring exon, could potentially increase the inclusion of the exon of interest as well.

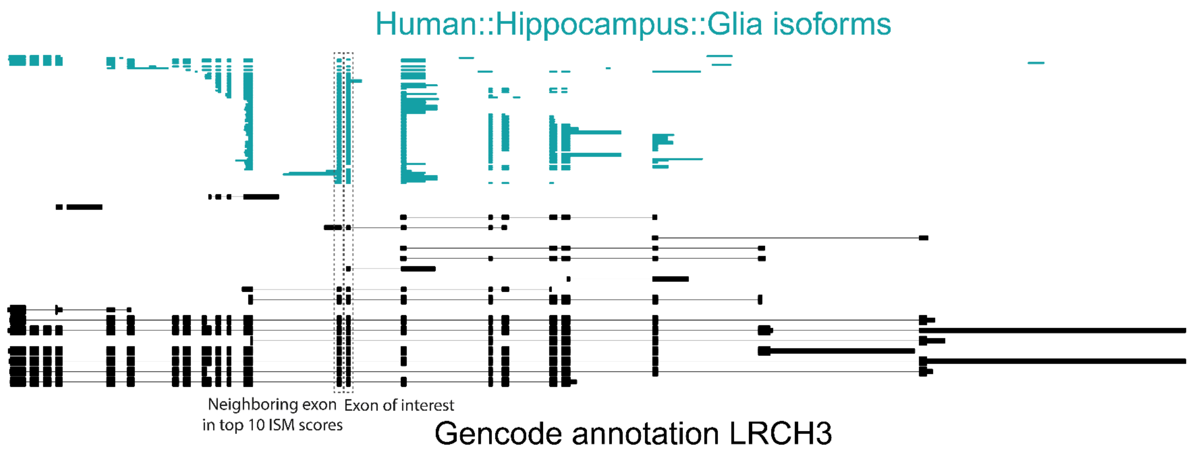


**Figure S19** Potential coordination in LRCH3. The exon of interest and the neighboring exon are either included together or not.

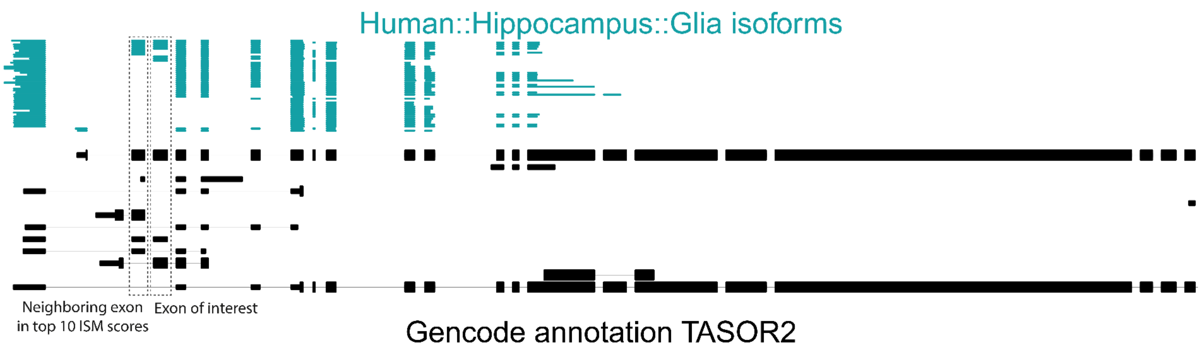


**Figure S20** Here it's random whether the neighboring exon and the exon of interest are included simultaneously, so no coordination.

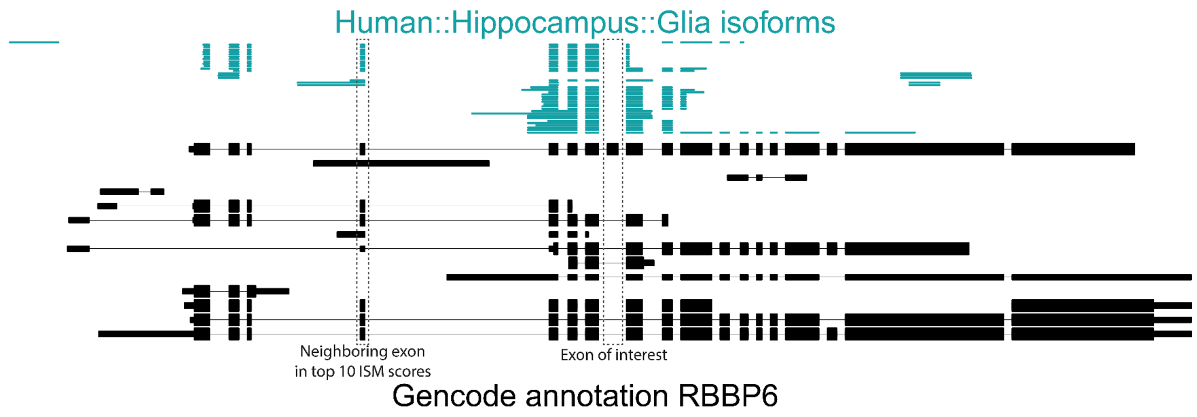


**Figure S21** The exon of interest is never included in our data in glia.

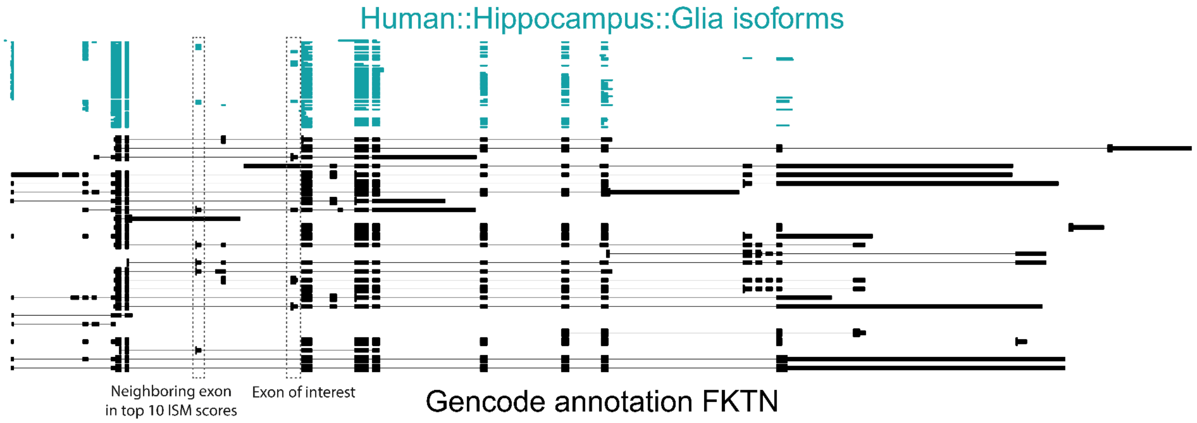


**Figure S22** Here it's random whether the neighboring exon and exon of interest are included separately or both.

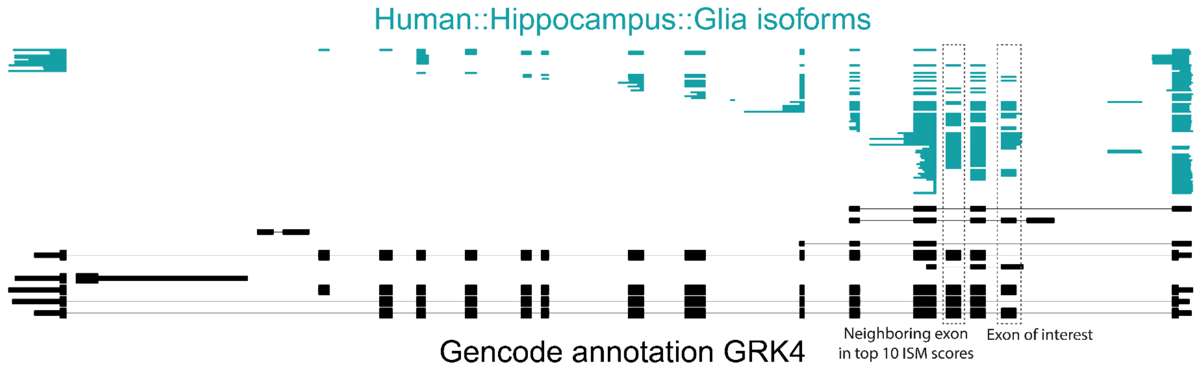


**Figure S23** Here it's random whether the neighboring exon and exon of interest are included separately or both.

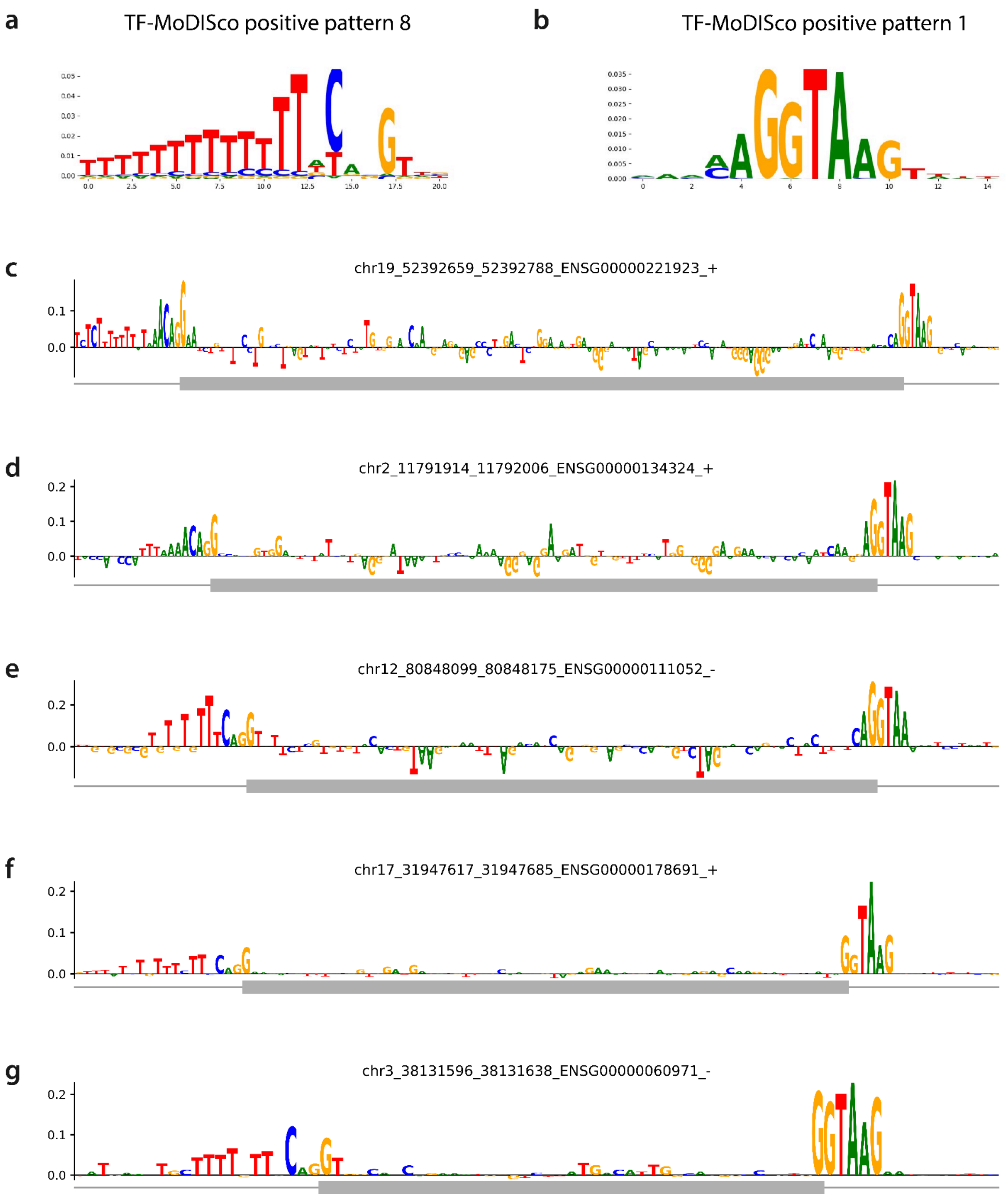


**Figure S24 a-b)** Example TF-MoDISco motifs that are related to the splice sites, **c-g)** ISM results of some sequences with these motifs.

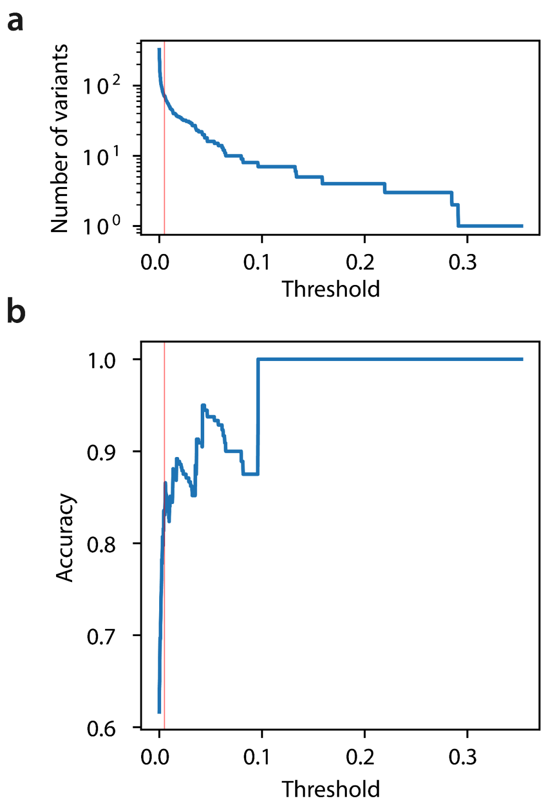


**Figure S25 a)** Number of variants for which the predicted effect is above the threshold, **b)** Accuracy for those variants.

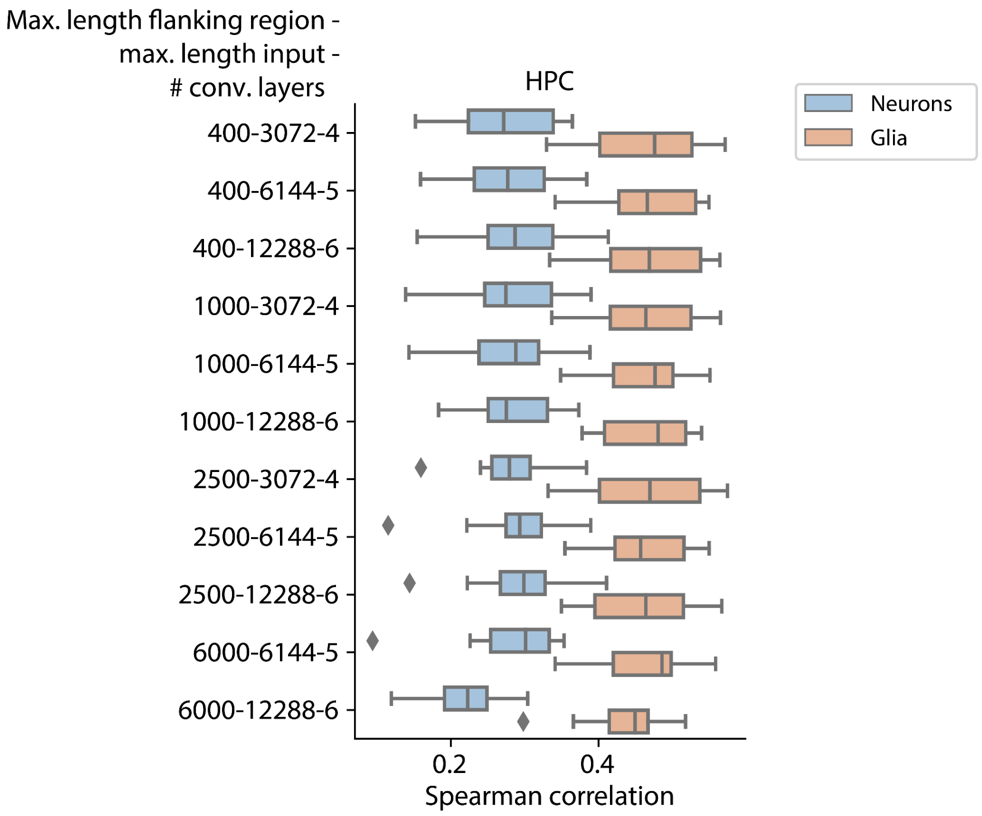


**Figure S26** Performance of the $DL_{all}$ models during the 10-fold cross-validation on variable exons from neurons and glia in the HPC. The models have different maximum lengths of flanking regions (so the sequence up- and downstream of the exon), maximum lengths of input sequences, and a different number of convolutional layers.

**Table S1** Number of exons in the training data for the different models. We split the training data into ten folds during the cross-validation. Exons are always in the same folds to allow for comparison across the models.

| | $DL_{all}$ | $LR_{all}$ | $DL_{var0.1}$ | $LR_{var0.1}$ | $LR_{var0.25}$ |
|---|---|---|---|---|---|
| HPC | 42,942 | 37,382 | 9,929 | 8,422 | 1,827 |
| FC | 15,747 | 14,077 | 2,637 | 2,315 | 802 |

**Table S2** Mouse data

| Brain region | Number of mice | Exons measured for both glia and neurons | Variable exons | Reference |
|---|---|---|---|---|
| HPC | 2 | 23,857 | 528 | Joglekar et al.[1] |
| VisC | 2 | 48,515 | 1,404 | Joglekar et al.[1] |

**Table S3** Differentially spliced RBPs

| Gene | Chr | Strand | Start | End | HPC - neur | HPC - glia |
|---|---|---|---|---|---|---|
| PABPC4 | 1 | - | 39563836 | 39563922 | 0.47 | 0.87 |
| | | | 39564686 | 39564773 | 0.66 | 0.92 |
| ZRANB2 | 1 | - | 71065678 | 71065752 | 0.64 | 0.33 |
| PUM2 | 2 | - | 20278583 | 20278819 | 0.22 | 0.36 |
| CPEB4 | 5 | + | 173943026 | 173943049 | 0.66 | 0.34 |
| DROSHA | 5 | - | 31521123 | 31521215 | 0.94 | 0.67 |
| XPO5 | 6 | - | 43549489 | 43549578 | 0.71 | 1.00 |
| | | | 43549893 | 43549934 | 0.72 | 1.00 |
| | | | 43558501 | 43558591 | 0.48 | 1.00 |
| TBRG4 | 7 | - | 45103333 | 45103443 | 0.94 | 0.64 |
| | | | 45104099 | 45104256 | 0.83 | 0.39 |
| | | | 45104538 | 45104709 | 0.67 | 0.40 |
| | | | 45105441 | 45105764 | 0.93 | 0.64 |
| YBX3 | 12 | - | 10709908 | 10710114 | 0.92 | 0.57 |
| PUS1 | 12 | + | 131939173 | 131939275 | 0.98 | 0.71 |
| TAF15 | 17 | + | 35833907 | 35833941 | 0.63 | 0.90 |
| EFTUD2 | 17 | - | 44854556 | 44854682 | 0.56 | 0.93 |
| | | | 44854918 | 44855004 | 0.50 | 0.92 |
| | | | 44857075 | 44857157 | 0.68 | 0.94 |
| | | | 44859905 | 44860045 | 0.63 | 0.90 |
| | | | 44863655 | 44863782 | 0.47 | 0.92 |
| FXR2 | 17 | - | 7594238 | 7594347 | 0.69 | 1.00 |

**Table S4** Mapping of exon names to genomic coordinates. We used reference genome GRCh38 to get the genomic coordinates and GENCODE v35 to count exons.

| **Gene name** | **Chr** | **Strand** | **Exon ID** | **Start** | **End** |
|---|---:|---|---|---:|---:|
| *XRN2* | 20 | + | Exon 21 | 21354789 | 21354872 |
| | | | Exon 22 | 21356080 | 21356177 |
| | | | Exon 24 | 21357736 | 21657792 |
| *TPCN1* | 12 | + | Exon 42 | 113278189 | 113278237 |
| *ZNF880* | 19 | + | Exon 15 | 52383665 | 52383764 |
| *RARS1* | 5 | + | Exon 25 | 168497947 | 168498046 |
