## Supplementary File 1 for "Predicting cell-type-specific exon inclusion in the human brain reveals more complex splicing mechanisms in neurons than glia"

| pattern | num_seqlets | modisco_cwm_fwd | modisco_cwm_rev | match0 | qval0 | match0_logo | match1 | qval1 | match1_logo | match2 | qval2 | match2_logo |
|---|---|---|---|---|---|---|---|---|---|---|---|---|
| pos_patterns.pattern_0 | 2806 | | | TRA2A_7mer_logo0 | 1.000000 | | PABPN1_7mer_logoM148 | 1.000000 | | tra2_7mer_logoM076 | 1.000000 | |
| pos_patterns.pattern_1 | 1415 | | | PCBP1_7mer_logoM177 | 1.000000 | | DAZAP1_7mer_logoM013 | 1.000000 | | Hrb27C_7mer_logoM028 | 1.000000 | |
| pos_patterns.pattern_2 | 1120 | | | TVAG514790_7mer_logoM204 | 0.012887 | | PABPC4_7mer_logoM042 | 0.012887 | | gw184451_7mer_logoM198 | 0.016268 | |
| pos_patterns.pattern_3 | 983 | | | HNRNPC_7mer_logo0 | 0.025064 | | Sxl_7mer_logoM114 | 0.031924 | | gw184451_7mer_logoM198 | 0.046902 | |
| pos_patterns.pattern_4 | 977 | | | RBM45_7mer_logo5 | 0.413316 | | Rsf1_7mer_logoM059 | 0.413316 | | CG2950_7mer_logoM007 | 1.000000 | |
| pos_patterns.pattern_5 | 901 | | | TAF15_7mer_logo0 | 0.012774 | | HNRNPA2B1_7mer_logo0 | 0.019606 | | SRSF8_7mer_logo1 | 0.019606 | |
| pos_patterns.pattern_6 | 640 | | | KHDRBS2_7mer_logo1 | 0.454750 | | KHDRBS3_7mer_logo0 | 0.454750 | | RBMS3_7mer_logo2 | 0.454750 | |
| pos_patterns.pattern_7 | 614 | | | IGF2BP1_7mer_logo1 | 0.793093 | | QKI_7mer_logoM046 | 0.793093 | | how_7mer_logoM113 | 0.793093 | |
| pos_patterns.pattern_8 | 547 | | | TRA2A_7mer_logo1 | 0.063024 | | PUF60_7mer_logo0 | 0.063024 | | PABPC4_7mer_logoM042 | 0.063024 | |
| pos_patterns.pattern_9 | 394 | | | B52_7mer_logoM126 | 0.022741 | | HNRNPF_7mer_logo1 | 0.022741 | | HNRNPH2_7mer_logoM151 | 0.022741 | |
| pos_patterns.pattern_10 | 360 | | | YB1_7mer_logoM111 | 0.852294 | | IGF2BP2_7mer_logo3 | 1.000000 | | YB1_7mer_logoM081 | 1.000000 | |
| pos_patterns.pattern_11 | 326 | | | Hrb87F_7mer_logoM029 | 0.005667 | | Hrb98DE_7mer_logoM092 | 0.005667 | | Hrb98DE_7mer_logoM093 | 0.005667 | |
| pos_patterns.pattern_12 | 276 | | | KHDRBS3_7mer_logo0 | 0.245746 | | RBMS3_7mer_logo2 | 0.245746 | | shep_7mer_logoM166 | 0.290158 | |
| pos_patterns.pattern_13 | 221 | | | RBM4B_7mer_logo2 | 1.000000 | | PCBP1_7mer_logo3 | 1.000000 | | RBM22_7mer_logo0 | 1.000000 | |
| pos_patterns.pattern_14 | 206 | | | SFPQ_7mer_logo0 | 0.737892 | | RBM45_7mer_logo2 | 0.737892 | | pum_7mer_logoM101 | 1.000000 | |
| pos_patterns.pattern_15 | 163 | | | Hrb98DE_7mer_logoM091 | 0.410690 | | HNRNPF_7mer_logo1 | 0.410690 | | DAZAP1_7mer_logoM013 | 0.410690 | |
| pos_patterns.pattern_16 | 143 | | | QKI_7mer_logoM046 | 0.063280 | | how_7mer_logoM113 | 0.063280 | | LmjF180180_7mer_logoM187 | 0.208011 | |
| pos_patterns.pattern_17 | 137 | | | RBM25_7mer_logo2 | 0.034396 | | PUM1_7mer_logo1 | 0.231354 | | EWSR1_7mer_logo0 | 0.247968 | |
| pos_patterns.pattern_18 | 132 | | | TAF15_7mer_logo1 | 0.590264 | | HNRNPH2_7mer_logoM151 | 0.590264 | | HNRNPF_7mer_logo0 | 0.909467 | |
| pos_patterns.pattern_19 | 128 | | | PCBP1_7mer_logo3 | 1.000000 | | PCBP4_7mer_logo2 | 1.000000 | | RBM22_7mer_logo0 | 1.000000 | |
| pos_patterns.pattern_20 | 111 | | | TAF15_7mer_logo0 | 0.107660 | | SRSF8_7mer_logo1 | 0.107660 | | RBM25_7mer_logo2 | 0.163125 | |
| pos_patterns.pattern_21 | 109 | | | NCU02404_7mer_logoM206 | 0.236015 | | tra2_7mer_logoM076 | 0.236015 | | SRSF1_7mer_logoM103 | 0.236015 | |

| pattern | num_seqlets | modisco_cwm_fwd | modisco_cwm_rev | match0 | qval0 | match0_logo | match1 | qval1 | match1_logo | match2 | qval2 | match2_logo |
|---|---|---|---|---|---|---|---|---|---|---|---|---|
| pos_patterns.pattern_22 | 87 | | | TRA2A_7mer_logo1 | 1.000000 | | TRA2A_7mer_logo0 | 1.000000 | | YBX2_7mer_logoM082 | 1.000000 | |
| pos_patterns.pattern_23 | 61 | | | SF1_7mer_logo2 | 1.000000 | | KHDRBS3_7mer_logo3 | 1.000000 | | ELAVL4_7mer_logo1 | 1.000000 | |
| pos_patterns.pattern_24 | 59 | | | RBM25_7mer_logo0 | 0.104614 | | HNRNPA2B1_7mer_logo0 | 0.104614 | | HNRNPH2_7mer_logoM151 | 0.104614 | |
| pos_patterns.pattern_25 | 55 | | | DAZAP1_7mer_logo2 | 0.940224 | | SNRPA_7mer_logoM348 | 0.940224 | | HNRNPA0_7mer_logo3 | 0.940224 | |
| pos_patterns.pattern_26 | 52 | | | PCBP1_7mer_logo1 | 0.425954 | | PCBP2_7mer_logo2 | 0.536159 | | PCBP4_7mer_logo1 | 0.536159 | |
| pos_patterns.pattern_27 | 51 | | | SAMD4A_7mer_logoM061 | 0.999996 | | RC3H1_7mer_logo1 | 0.999996 | | pUf68_7mer_logoM132 | 0.999996 | |
| pos_patterns.pattern_28 | 43 | | | TRNAU1AP_7mer_logo1 | 1.000000 | | BOLL_7mer_logo3 | 1.000000 | | MSI1_7mer_logo1 | 1.000000 | |
| pos_patterns.pattern_29 | 29 | | | tra2_7mer_logoM076 | 1.000000 | | TRA2A_7mer_logo0 | 1.000000 | | NaN | NaN | |
| pos_patterns.pattern_30 | 27 | | | EWSR1_7mer_logo3 | 1.000000 | | TRA2A_7mer_logo1 | 1.000000 | | SFPQ_7mer_logo0 | 1.000000 | |
| neg_patterns.pattern_0 | 1873 | | | NaN | NaN | | NaN | NaN | | NaN | NaN | |
| neg_patterns.pattern_1 | 915 | | | HNRNPH2_7mer_logoM151 | 0.010881 | | HNRNPF_7mer_logo0 | 0.016703 | | RBM23_7mer_logo0 | 0.016703 | |
| neg_patterns.pattern_2 | 883 | | | TAF15_7mer_logo0 | 0.006430 | | PCBP1_7mer_logo0 | 0.016483 | | SRSF8_7mer_logo1 | 0.016483 | |
| neg_patterns.pattern_3 | 743 | | | NaN | NaN | | NaN | NaN | | NaN | NaN | |
| neg_patterns.pattern_4 | 699 | | | Tbg97295210_7mer_logoM216 | 0.008832 | | Fusip1_7mer_logoM087 | 0.017973 | | PUF60_7mer_logo0 | 0.017973 | |
| neg_patterns.pattern_5 | 628 | | | Sxl_7mer_logoM114 | 0.018752 | | TVAG514790_7mer_logoM204 | 0.018752 | | PABPC4_7mer_logoM042 | 0.018752 | |
| neg_patterns.pattern_6 | 540 | | | PTBP3_7mer_logo0 | 1.000000 | | KHDRBS3_7mer_logo3 | 1.000000 | | PTBP3_7mer_logo1 | 1.000000 | |
| neg_patterns.pattern_7 | 540 | | | HNRNPH2_7mer_logoM151 | 0.014835 | | HNRNPF_7mer_logo0 | 0.014835 | | HNRNPF_7mer_logo1 | 0.014835 | |
| neg_patterns.pattern_8 | 400 | | | TRNAU1AP_7mer_logo2 | 0.592513 | | KHDRBS3_7mer_logo0 | 0.592513 | | RBM15B_7mer_logo1 | 0.592513 | |
| neg_patterns.pattern_9 | 386 | | | PUF60_7mer_logo0 | 0.060297 | | HNRNPC_7mer_logo0 | 0.060297 | | PABPC4_7mer_logoM042 | 0.060297 | |
| neg_patterns.pattern_10 | 358 | | | TAF15_7mer_logo0 | 0.217791 | | SRSF8_7mer_logo1 | 0.217791 | | SFPQ_7mer_logo3 | 0.217791 | |
| neg_patterns.pattern_11 | 311 | | | HNRNPC_7mer_logo0 | 0.040238 | | Sxl_7mer_logoM114 | 0.040438 | | ELAVL1_7mer_logoM232 | 0.040438 | |
| neg_patterns.pattern_12 | 295 | | | CPEB1_7mer_logo0 | 0.671904 | | DAZ3_7mer_logo3 | 0.671904 | | KHDRBS2_7mer_logo1 | 0.671904 | |

| pattern | num_seqlets | modisco_cwm_fwd | modisco_cwm_rev | match0 | qval0 | match0_logo | match1 | qval1 | match1_logo | match2 | qval2 | match2_logo |
|---|---|---|---|---|---|---|---|---|---|---|---|---|
| neg_patterns.pattern_13 | 292 | | | CG2931_7mer_logoM138 | 0.109767 | | BOLL_7mer_logo2 | 0.125769 | | RBM42_7mer_logoM142 | 0.125769 | |
| neg_patterns.pattern_14 | 269 | | | HNRNPC_7mer_logo0 | 0.016414 | | Sxl_7mer_logoM114 | 0.016414 | | TVAG514790_7mer_logoM204 | 0.016414 | |
| neg_patterns.pattern_15 | 243 | | | HNRNPF_7mer_logo1 | 1.000000 | | EWSR1_7mer_logo3 | 1.000000 | | RBM25_7mer_logo2 | 1.000000 | |
| neg_patterns.pattern_16 | 227 | | | TIA1_7mer_logo0 | 0.034657 | | KHDRBS3_7mer_logo1 | 0.036074 | | ELAVL4_7mer_logo0 | 0.047855 | |
| neg_patterns.pattern_17 | 220 | | | exc7_7mer_logoM014 | 0.659310 | | ELAVL4_7mer_logo2 | 0.820726 | | TRA2A_7mer_logo0 | 0.820726 | |
| neg_patterns.pattern_18 | 204 | | | HNRNPA2B1_7mer_logo0 | 0.018745 | | EWSR1_7mer_logo3 | 0.018745 | | HNRNPH2_7mer_logoM151 | 0.018745 | |
| neg_patterns.pattern_19 | 193 | | | shep_7mer_logoM066 | 0.454675 | | KHDRBS2_7mer_logo0 | 0.454675 | | shep_7mer_logoM166 | 0.454675 | |
| neg_patterns.pattern_20 | 192 | | | A1CF_7mer_logo0 | 0.237324 | | TRNAU1AP_7mer_logo1 | 0.237324 | | HNRNPD_7mer_logo0 | 0.237324 | |
| neg_patterns.pattern_21 | 178 | | | RBM25_7mer_logo1 | 0.985434 | | PCBP2_7mer_logo1 | 0.985434 | | RBM23_7mer_logo0 | 0.985434 | |
| neg_patterns.pattern_22 | 146 | | | KHDRBS3_7mer_logo0 | 1.000000 | | RBMS3_7mer_logo2 | 1.000000 | | CELF1_7mer_logo2 | 1.000000 | |
| neg_patterns.pattern_23 | 140 | | | ELAVL1_7mer_logoM232 | 0.001467 | | gw184451_7mer_logoM198 | 0.001467 | | HNRNPC_7mer_logo0 | 0.008244 | |
| neg_patterns.pattern_24 | 139 | | | tra2_7mer_logoM076 | 0.080765 | | TRA2A_7mer_logo0 | 0.097847 | | shep_7mer_logoM165 | 0.760849 | |
| neg_patterns.pattern_25 | 128 | | | PPRC1_7mer_logoM044 | 0.000002 | | RBM4_7mer_logoM050 | 0.005436 | | RBM4B_7mer_logo0 | 0.005436 | |
| neg_patterns.pattern_26 | 120 | | | MSI1_7mer_logoM040 | 1.000000 | | MSI1_7mer_logoM167 | 1.000000 | | RBM25_7mer_logo2 | 1.000000 | |
| neg_patterns.pattern_27 | 100 | | | SRSF5_7mer_logo2 | 0.637969 | | TAF15_7mer_logo1 | 0.637969 | | PCBP2_7mer_logo0 | 0.637969 | |
| neg_patterns.pattern_28 | 94 | | | SFPQ_7mer_logo4 | 0.205755 | | CNOT4_7mer_logo2 | 0.205755 | | PCBP1_7mer_logo0 | 0.205755 | |
| neg_patterns.pattern_29 | 71 | | | EIF4G2_7mer_logo3 | 0.031424 | | PPRC1_7mer_logoM044 | 0.035493 | | SFPQ_7mer_logo4 | 0.035493 | |
| neg_patterns.pattern_30 | 62 | | | RBM41_7mer_logoM051 | 1.000000 | | RC3H1_7mer_logo0 | 1.000000 | | shep_7mer_logoM165 | 1.000000 | |
| neg_patterns.pattern_31 | 61 | | | FMR1_7mer_logoM016 | 0.366338 | | FXR2_7mer_logoM020 | 0.366338 | | SRSF2_7mer_logo3 | 0.952385 | |
| neg_patterns.pattern_32 | 60 | | | CG2931_7mer_logoM138 | 0.158612 | | RBM42_7mer_logoM142 | 0.315635 | | BOLL_7mer_logo2 | 1.000000 | |
| neg_patterns.pattern_33 | 60 | | | PPRC1_7mer_logoM044 | 0.033704 | | SRSF5_7mer_logo2 | 0.033704 | | TAF15_7mer_logo2 | 0.033704 | |
| neg_patterns.pattern_34 | 57 | | | Hrb27C_7mer_logoM090 | 0.287879 | | Hrb27C_7mer_logoM028 | 0.287879 | | ANIA04546_7mer_logoM226 | 0.287879 | |

| pattern | num_seqlets | modisco_cwm_fwd | modisco_cwm_rev | match0 | qval0 | match0_logo | match1 | qval1 | match1_logo | match2 | qval2 | match2_logo |
|---|---|---|---|---|---|---|---|---|---|---|---|---|
| neg_patterns.pattern_35 | 52 | | 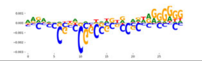 | PCBP1_7mer_logo0 | 0.215737 | 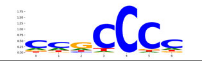 | PPRC1_7mer_logoM044 | 0.215737 | 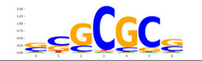 | TAF15_7mer_logo1 | 0.215737 | 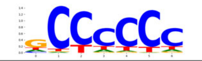 |
| neg_patterns.pattern_36 | 48 | 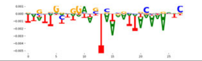 | | TRNAU1AP_7mer_logo1 | 0.348772 | 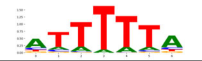 | NUPL2_7mer_logo0 | 0.657776 | 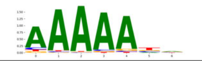 | TRNAU1AP_7mer_logo0 | 0.657776 | 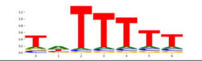 |
| neg_patterns.pattern_37 | 45 | 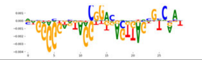 | | RBM6_7mer_logo3 | 1.000000 | 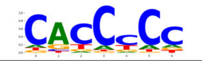 | SRSF5_7mer_logo4 | 1.000000 | 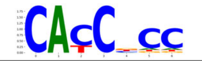 | NaN | NaN | |
| neg_patterns.pattern_38 | 36 | 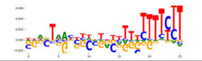 | | SRSF5_7mer_logo4 | 0.349634 | 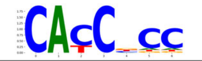 | tra2_7mer_logoM076 | 0.349634 | 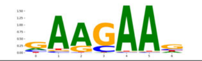 | PTBP1_7mer_logoM227 | 0.349634 | 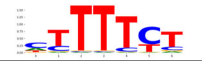 |
| neg_patterns.pattern_39 | 35 | 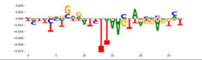 | | exc7_7mer_logoM014 | 0.996656 | 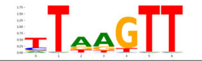 | DAZAP1_7mer_logo2 | 0.996656 | 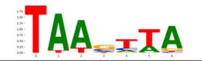 | ELAVL4_7mer_logo2 | 1.000000 | 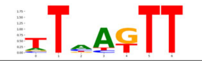 |
| neg_patterns.pattern_41 | 30 | 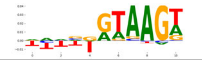 | | SFPQ_7mer_logo0 | 0.321676 | 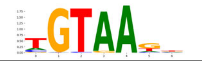 | QKI_7mer_logoM046 | 0.613013 | 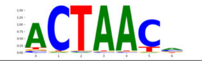 | how_7mer_logoM113 | 0.613013 | 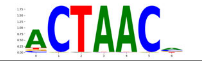 |
| neg_patterns.pattern_43 | 26 | 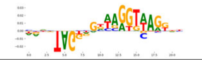 | | NCU02404_7mer_logoM206 | 0.565431 | 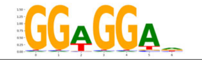 | HNRNPF_7mer_logo2 | 0.565431 | 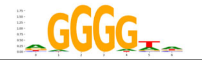 | LIN28A_7mer_logoM035 | 0.565431 | 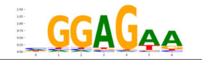 |
